# Identification of Promoter Activity in Gene-Less Cassettes from *Vibrionaceae* Superintegrons

**DOI:** 10.1101/2023.11.21.568050

**Authors:** Paula Blanco, Alberto Hipólito, Lucía García-Pastor, Filipa Trigo da Roza, Laura Toribio-Celestino, Alba Cristina Ortega, Ester Vergara, Álvaro San Millán, José Antonio Escudero

## Abstract

Integrons are genetic platforms that acquire new genes encoded in integron cassettes (ICs), building arrays of adaptive functions. ICs generally encode promoterless genes, whose expression relies on the platform-associated Pc promoter, with the cassette array functioning as an operon-like structure regulated by the distance to the Pc. This is relevant in large sedentary chromosomal integrons (SCIs) carrying hundreds of ICs, like those in *Vibrio* species. We selected 29 gene-less cassettes in four *Vibrio* SCIs, and explored whether their function could be related to the transcription regulation of adjacent ICs. We show that most gene-less cassettes have promoter activity on the sense strand, enhancing the expression of downstream cassettes. Additionally, we identified the transcription start sites of gene-less ICs through 5’-RACE. Accordingly, we found that most of the superintegron in *Vibrio cholerae* is not silent. These *promoter cassettes* can trigger the expression of a silent *dfrB9* cassette downstream, increasing trimethoprim resistance >512-fold in *V. cholerae* and *Escherichia coli*. Furthermore, one cassette with an antisense promoter can reduce trimethoprim resistance when cloned downstream. Our findings highlight the regulatory role of gene-less cassettes in the expression of adjacent cassettes, emphasizing their significance in SCIs and their clinical importance if captured by mobile integrons.

## INTRODUCTION

Integrons are bacterial genetic elements that capture and rearrange genes embedded in structures known as integron cassettes (ICs) (reviewed in ^1–3^). Integrons are known for their key role in the rise and spread of antibiotic resistance (AR) genes. Carried on conjugative plasmids, <u>m</u>obile integrons (MIs) circulate among the most important Gram-negative pathogens, disseminating more than 170 resistance determinants against most antibiotics ^4^. Yet, integrons are naturally found in almost 20% of bacterial genomes available in the databases. These are usually referred to as *sedentary chromosomal integrons* (SCIs) to distinguish them from MIs ^5^. Among SCIs, those in *Vibrio* species are archetypical, with very large arrays that reach 300 cassettes. Indeed, *V. cholerae*‘s superintegron was the first SCI discovered and is today the most studied and the paradigm in the field. Both types of integrons are not isolated form one another: mobile integrons are massively shed to the environment in the faeces of humans and animals ^6^. The conjugative transfer to and from environmental bacteria allows MIs to capture cassettes from SCIs ^7^ and to bring them back to clinical settings. There, they can easily spread if they provide an adaptive advantage, as is the case of AR genes.

All integrons are composed of a stable and a variable region. The stable region is a genetic platform that contains the integrase-coding gene (*intI*) with its associated promoter (Pint), the *attI* recombination site where integron cassettes are inserted, and the constitutive promoter PC, that drives their transcription. The variable region of any integron is the cassette array, that can be very different between strains and represents a low-cost memory of valuable functions for the cell (Figure 1) ^8^.

**Figure 1.**
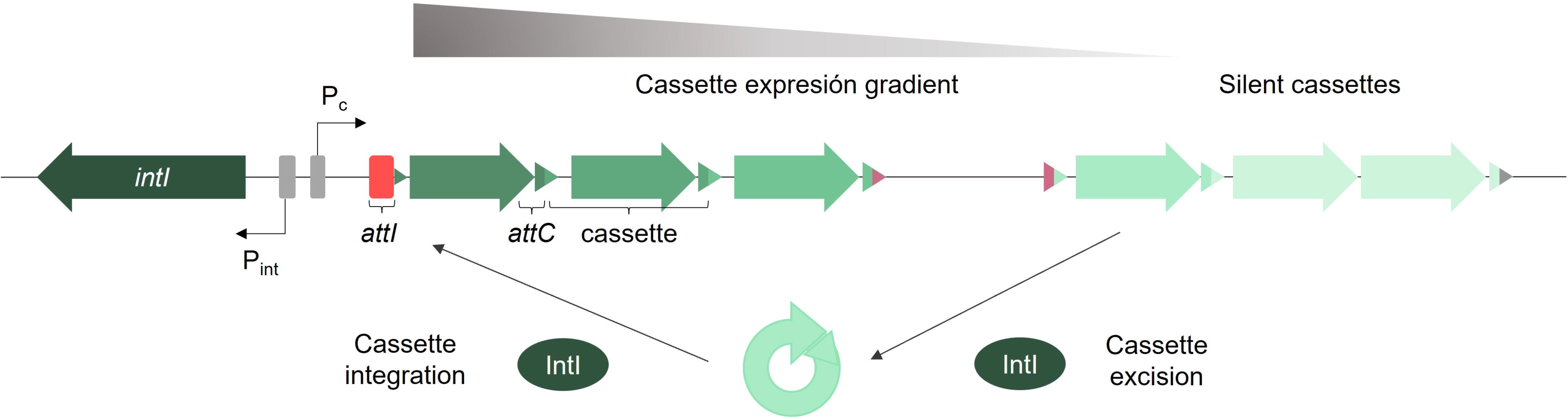
Typical structure of an integron. Integrons are composed of the integrase-coding gene (*intI*) and its promoter (Pint), an integron recombination site (*attI*), and a variable cassette array whose expression is driven by the PC promoter. Genes are depicted as arrows. Cassette recombination sites (*attC*) are depicted as triangles. Garnet *attC* sites delimit a gene-less cassette. Integron cassettes can be integrated and excised from the platform through integrase-mediated recombination reactions between *attI* and *attC* sites. This can lead to the reshuffling of cassettes and the modulation of their expression.

Cassettes typically consist of a single promoterless gene and a recombination site, named *attC* ^9,10^. The integron integrase captures cassettes through site-specific recombination between the *attI* site in the platform and the *attC* in the cassette ^11^. Once acquired, cassettes are rendered functional by the activity of the PC. Subsequent acquisition events at *attI* push existing cassettes away from the PC decreasing their expression. Nevertheless, cassettes in the array can be excised through the recombination of contiguous *attC* sites ^12^ and reintegrated in first position, ultimately increasing their expression levels ^13^. The lack of promoters in cassettes and the reshuffling of the array, supports the role of the PC as the exclusive promoter of the array. However, some exceptions to this rule have also been reported. Internal promoters have been identified in several ICs, such as toxin-antitoxin systems, the erythromycin resistance gene *ere(A)*, the chloramphenicol resistance gene *cmlA*, or the quinolone resistance gene *qnrVC1* ^14–16^. Yet, the impact that such promoters can have on downstream genes within arrays remains today largely unexplored.

Although ICs generally contain a single ORF that occupies most of the cassette, some studies have reported the presence of cassettes that do not contain such ORFs, and hence seem to have a biological role other than encoding a protein. It has been proposed that these gene-less cassettes may act as regulatory elements within an integron, including features like binding sites for regulatory proteins, small RNAs, or promoters, facilitating the evolution of genomes ^17,18^. The presence of internal promoters in gene-less cassettes has actually been experimentally demonstrated, but only in two cassettes from the atypical *Treponema* integrons ^19^.

In this study, we tested whether gene-less cassettes act as internal promoters within cassette arrays in SCIs in *Vibrionaceae*. To that end, we scanned the SCIs of four *Vibrionaceae* family species (*V. cholerae*, *V. fischeri*, *V. parahaemolyticus* and *V. vulnificus*) looking for gene-less cassettes. We identified and synthesized 29 gene-less cassettes and cloned all of them on a bidirectional reporter plasmid (pDProm), based on the expression of *mCherry* and *gfp*, and quantitated promoter activity on both DNA strands by fluorescence intensity. We found that promoter activity was common among cassettes in both strands, although with a clear bias towards the sense strand. Notably, certain gene-less cassettes led to a more than 100-fold increase in GFP fluorescence, suggesting that expression of cassettes can be strongly modulated by gene-less cassettes located upstream. We further corroborated these findings by confirming promoter activity through RT-qPCR. Additionally, we described the transcription star sites of a selection of ICs by performing 5’-RACE. Replacing the *gfp* gene for a trimethoprim resistance cassette (*dfrB9*), led us to find that at least two gene-less cassettes conferred a clinically relevant increase in trimethoprim resistance, both in *V. cholerae* and *E. coli*. Furthermore, we uncovered that the presence of transcription on the antisense strand within a cassette could effectively silence an upstream *dfrB9* cassette, leading to a reduction in antibiotic resistance levels. We therefore reveal that gene-less cassettes from large SCIs play a regulatory role promoting the expression of distantly located cassettes, independent of their proximity to the PC promoter. These ICs can have a strong impact in AR if integrated into mobile integrons.

## MATERIALS AND METHODS

### Bacterial strains, plasmids, and culture conditions

All bacterial strains and plasmids used in this study are listed in Table 1. All strains were cultured in both solid or liquid Luria-Bertani (LB) medium (BD) at 37°C. Kanamycin (Sigma Aldrich) was added when required at the following concentrations: 25 µg/mL for *E. coli* and 75 µg/mL for *V. cholerae*.

**Table 1.** Strains used in this study.

| Strains |  | Plasmid/Cassette cloned in<br>pDProm | Origin species | Genome |
| --- | --- | --- | --- | --- |
| *E. coli*<br>DH5α | *V.*<br>*cholerae*<br>N16961 |  |  |  |
| Lab strain | - | - | - | - |
| - | 59 | - | - | - |
| A369 | - | pSU38 P<sub>lac</sub>::*gfp* (pA369) | - | - |
| A866 | B118 | pDProm empty vector | - | - |
| A371 | B950 | pSU38 P<sub>CW</sub>::*gfp* | - | - |
| A370 | B949 | pSU38 P<sub>CS</sub>::*gfp* | - | - |
| A887 | B014 | P<sub>lac</sub> *attC gfp* | - | - |
| A888 | B012 | P<sub>lac</sub> *attC mCherry* | - | - |
| B022 | B103 | C1 (VCA0294-VCA0297) | *V. cholerae* N16961 | NZ_CP028828.1 |
| B023 | B104 | C2 (VCA0325-VCA0328) | *V. cholerae* N16961 | NZ_CP028828.1 |
| A998 | B078 | C3 (VCA0382a-VCA0387) | *V. cholerae* N16961 | NZ_CP028828.1 |
| B024 | B106 | C4 (VCA0389-VCA0391) | *V. cholerae* N16961 | NZ_CP028828.1 |
| A999 | B079 | C5 (VCA0392-VCA0395) | *V. cholerae* N16961 | NZ_CP028828.1 |
| A983 | B080 | C6 (VCA0403-VCA0405) | *V. cholerae* N16961 | NZ_CP028828.1 |
| A984 | B107 | C7 (VCA0428-VCA0431) | *V. cholerae* N16961 | NZ_CP028828.1 |
| B025 | B108 | C8 (VCA0436-VCA0439) | *V. cholerae* N16961 | NZ_CP028828.1 |
| B066 | B109 | C9 (VCA0460-VCA0463) | *V. cholerae* N16961 | NZ_CP028828.1 |
| B007 | B077 | C10 (VCA0498-VCA0501) | *V. cholerae* N16961 | NZ_CP028828.1 |
| A977 | B016 | F1 (VF_A0658- VF_A0659) | *V. fischeri* ES114 | NC_006841 |
| A978 | B017 | F2 (VF_A0655- VF_A0656) | *V. fischeri* ES114 | NC_006841 |
| A893 | B020 | F3 (VF_A0636- VF_A0637) | *V. fischeri* ES114 | NC_006841 |
| B072 | B110 | P1 (D0853_RS06935-D0853_RS06955) | *V. parahaemolyticus*<br>VPD14 | NZ_CP031781 |
| A953 | B021 | P2 (D0853_RS07000-D0853_RS07035) | *V. parahaemolyticus*<br>VPD14 | NZ_CP031781 |
| B073 | B111 | P3 (D0853_RS07065-D0853_RS07090) | *V. parahaemolyticus*<br>VPD14 | NZ_CP031781 |
| A997 | B081 | P4 (D0853_RS07095-D0853_RS07105) | *V. parahaemolyticus*<br>VPD14 | NZ_CP031781 |
| B074 | B112 | V1 (FOB71_ RS17245-FOB71_ RS17270) | *V. vulnificus*<br>FDAARGOS_663 | NZ_CP044069 |
| B076 | B114 | V2 (FOB71_ RS17150-FOB71_ RS17175) | *V. vulnificus*<br>FDAARGOS_663 | NZ_CP044069 |
| B069 | B115 | V3 (FOB71_ RS17135-FOB71_ RS17150) | *V. vulnificus*<br>FDAARGOS_663 | NZ_CP044069 |
| B070 | B116 | V4 (FOB71_ RS167700-FOB71_ RS16720) | *V. vulnificus*<br>FDAARGOS_663 | NZ_CP044069 |
| A985 | B082 | V5 (FOB71_ RS167700-FOB71_ RS16720) | *V. vulnificus*<br>FDAARGOS_663 | NZ_CP044069 |
| A986 | B083 | V6 (FOB71_ RS16665-FOB71_ RS16680) | *V. vulnificus*<br>FDAARGOS_663 | NZ_CP044069 |
| A987 | B084 | V7 (FOB71_RS16650-FOB71_RS16665) | *V. vulnificus*<br>FDAARGOS_663 | NZ_CP044069 |
| A988 | B085 | V8 (FOB71_RS16635-FOB71_RS16650) | *V. vulnificus*<br>FDAARGOS_663 | NZ_CP044069 |
| A989 | B086 | V9 (FOB71_RS16445-FOB71_RS16460) | *V. vulnificus*<br>FDAARGOS_663 | NZ_CP044069 |
| A990 | B087 | V10 (FOB71_RS16125-FOB71_RS1645) | *V. vulnificus*<br>FDAARGOS_663 | NZ_CP044069 |
| A991 | B088 | V11 (FOB71_RS16125-FOB71_RS1645) | *V. vulnificus*<br>FDAARGOS_663 | NZ_CP044069 |
| A992 | B089 | V12 (FOB71_RS15850-FOB71_RS15865) | *V. vulnificus*<br>FDAARGOS_663 | NZ_CP044069 |
| B071 | B117 | V13 (FOB71_RS15650-FOB71_RS15665) | *V. vulnificus*<br>FDAARGOS_663 | NZ_CP044069 |
| C435 | C438 | 450-bp *lacZ* fragment | *V. cholerae* N16961 | NZ_CP028827.1 |
| C436 | C439 | 550-bp *lacZ* fragment | *V. cholerae* N16961 | NZ_CP028827.1 |
| C437 | C440 | 650-bp *lacZ* fragment | *V. cholerae* N16961 | NZ_CP028827.1 |
| B933 | B909 | *dfrB9* | *Enterobacter cloacae* p068-50543A | ID: KC675185.1 |
| B895 | B897 | *P<sub>lac</sub>-dfrB9* | - | - |
| B936 | B912 | *C7-dfrB9* | - | - |
| B937 | B913 | *C8-dfrB9* | - | - |
| B935 | B911 | *F3-dfrB9* | - | - |
| B934 | B910 | *P4-dfrB9* | - | - |
| B938 | B914 | *V13-dfrB9* | - | - |
| C266 | C268 | *C8-dfrB9-F1* | - | - |
| C267 | C269 | *C8-dfrB9-C7* | - | - |

### Gene-less cassettes fragments design

An *in silico* analysis using Geneious Prime software (version 2.2) was carried out for the identification of ICs of several *Vibrionaceae* species that do not contain annotations nor potential genes (i.e.: ORFs smaller than 210bp (70 amino acids)) beginning with ATG, TTG or GTG ^20^: nine gene-less cassettes were identified in *V. cholerae* N16961 (NZ_CP028828.1); three in *V. fischeri* ES114 (NC_006841); four in *V. parahaemolyticus* VPD14 (NZ_CP031781); and thirteen in *V. vulnificus* FDAARGOS_663 (NZ_CP044069). DNA fragments were designed for synthesis by adding to each gene-less cassette fragment its own *attC* site and approximately 20 bp homology upstream and downstream of the pDProm plasmid backbone (Supplementary Figure S1). An ORF-containing cassette (78 amino acids) with no gene annotation, was also selected from the superintegron of *V. cholerae* N16961 (cassette C7) as an ORF-carrying cassette control. It is of note that *attC* sites are composed of R’’, L’’, L’ and R’ boxes (Supplementary Figure S1). The consensus sequence of the R’ box is 5’-G^↓^TTRRRY-3’, and it is where the crossover takes place (arrow). Hence, when a cassette is incorporated to the array, the R’ box is split in two parts: the RRRY nucleotides are at the 5’end of the cassette, fused to the *attC* of the previous cassette. This means that, when looking at cassette arrays, the exact bases in R’ box are not conserved, but rather contingent on the identity of the cassette downstream. When designing the cloning strategy of cassettes, we could not rule out this fact could have an influence in our results. We hence sought to add a consensus RRRY sequence to the R’ box of the *attC* sites of our cassettes. To do so, we extracted and aligned the RRRY bases at the 3’ of all selected cassettes. The nucleotides ATGT were identified as the most frequent in this subset of *attCs* (Supplementary Figure S1, inset). Despite not being exactly RRRY (R is G or A but not T), this sequence was fused in all cassettes to the conserved GTT triplet, conforming a consensus R’ box that is biologically representative. We then added approximately 20 bp homology with the plasmid backbone upstream and downstream the synthetic fragments and cloned them into pDProm using Gibson assembly. Additionally, two control fragments were designed containing either *gfp* or *mCherry* under the control of a Plac promoter upstream of a *V. cholerae* 128 bp-*attC* site (VCA0398) to detect possible effects that these recombination regions may have on transcription. The control fragments also contain approximately 20 bp homology with the plasmid backbone upstream and downstream for further cloning. The designed DNA fragments containing the gene-less cassettes and the control cassettes were synthesized by Integrated DNA Technologies (IDT). Since gene-less cassettes do not contain any annotations, they are named in this work with a letter belonging to the species they belong to, followed by correlative numbering. In Table 1, the cassettes upstream and downstream the gene-less cassettes are specified. Gene-less cassettes sequences and control fragments are listed in Supplementary Table S1.

### Construction of the plasmid pDProm and derivatives

Primers used in this study are described in Supplementary Table S2. The mCherry-coding gene was synthesised by IDT with 20 bp homology, upstream and downstream, with the pA369 plasmid, a pSU38 that contains the *gfp* gene (Lab collection). The pA369 backbone was amplified by PCR using primers bb_pSU38_long F and bb_pSU38_long R. The *mCherry*-containing fragment was cloned into the pA369 amplified backbone using Gibson assembly ^21^, giving rise to the plasmid pDProm. Briefly, a 4 µL reaction is prepared by mixing 1 µL of insert, 1 µL of linearized plasmid and 2 µL of the 2X Gibson assembly buffer (5X ISOBuffer, 10,000 u/ml T5 exonuclease, 2,000 u/ml Phusion polymerase, 40,000 u/ml Taq ligase, dH2O). The mix is incubated for 30 min at 50°C. The Gibson assembly product was then transformed into *E. coli* DH5α competent cells. Then, pDProm was extracted with the GeneJET Plasmid Miniprep (Thermo Scientific), according to the manufacturer’s instructions.

To evaluate the promoter activity of the selected cassettes, the pDProm backbone was amplified by PCR using primers bb_pSU38_F and bb_pSU38_mCherry_R. Subsequently, all synthesized cassettes-containing fragments were cloned between the two reporter genes following the previously described Gibson assembly protocol. As an additional control, three fragments derived from the *lacZ* gene (VC00556) of varying sizes (450-, 550-, and 650 bp) were cloned into pDProm. The amplification of the three fragments was performed with primers pDProm_lacZ_F, and pDProm_lacZ450_R, pDProm_lacZ550_R, or pDProm_lacZ650_R, respectively. Gibson assembly was carried out for the cloning of the three *lacZ* fragments into pDProm. The resulting pDProm derivatives were then transformed into *E. coli* DH5α competent cells. The presence of the inserts was confirmed by PCR using primers MRVII and MFD and verified by DNA sequencing.

### Transformation of pDProm plasmids in *V. cholerae*

All pDProm plasmids, containing the gene-less cassettes or *lacZ* fragments as inserts, were extracted from *E. coli* DH5α cultures using the GeneJET Plasmid Miniprep (Thermo Scientific), following the manufacturer’s instructions. For the preparation of *V. cholerae* electrocompetent cells, a single colony of *V. cholerae* N16961 was initially inoculated in 3 mL of LB medium and cultured for 20 h at 37°C. Then, a 1:100 dilution was performed in 10 mL of LB and cultures were grown until reaching an OD600 of 0,5. The cultures were washed three times in G buffer (137 mM sucrose, 1 mM HEPES, pH 8.0) by centrifuging at 4,000 rpm and 4°C for 8 min. The final pellet was resuspended in 80 µL of G buffer.

Electrocompetent cells (40 µL) were mixed with 1 µL of each purified plasmid in a prechilled Eppendorf tube and transferred to a 0,1-cm electroporation cuvette (MBP). Electroporation was performed at 1,400 V for a 5 ms pulse using the Eporator (Eppendorf). Sequentially, cells were cultured with 1 mL of LB for 1 h at 37°C. Selection of *V. cholerae* clones containing the pDProm and derivative plasmids was accomplished by adding kanamycin (75 µg/mL). Plasmids presence was confirmed by PCR with primers MRVII and MFD and verified by DNA sequencing.

### Flow cytometry analysis

To determine promoter activity, bacterial strains carrying the pDProm plasmids were cultured on LB supplemented with kanamycin (75 µg/mL) for 20 h at 37°C. Samples were diluted 1:400 on filtered NaCl (0,9 M) in 96-well plates (Nunc, Thermo Scientific) and analysed with a CytoFLEX-S flow cytometer (Beckman Coulter). The 488- and the 661-nm lasers were used for GFP and mCherry excitation, respectively. GFP and mCherry fluorescence intensity was detected through 525/40 nm (FITC) and 610/20 nm (ECD) band pass filters, respectively. At least three biological replicates and two technical replicates were used, and 30,000 events per sample were recorded. Flow cytometry data were analysed with the CytExpert software (version 2.4; Beckman Coulter).

### RNA isolation and RNA-seq data analysis

Total RNA was extracted from *V. cholerae* N16961 cultures grown in LB at 37°C in both exponential (OD600 0,8) and stationary (OD600 2,8) growth phases using the RNeasy® Mini Kit (QIAGEN), following the manufacturer’s protocol. To eliminate any residual DNA, RNA was treated using the TURBO DNA-free™ Kit (Invitrogen). RNA concentration was measured using a BioSpectrometer (Eppendorf), and RNA integrity was determined using a Qubit™ 4 fluorometer (Invitrogen) with the RNA IQ Assay Kit (Invitrogen). Ribodepletion RNA library and sequencing was performed at the Oxford Genomics Centre using a NovaSeq6000 sequencing system (Illumina). Three biological samples per condition were sequenced. Additionally, RNA-Seq data of *V. cholerae* N16961 from an independent study (BioProject PRJNA420494; SRA identifiers: SRR6334020, SRR6334021 and SRR6334022) ^22^ was also analysed.

Raw reads from both studies were trimmed using Trim Galore v0.6.6 (https://github.com/FelixKrueger/TrimGalore) with a quality threshold of 20 and removing adapters and reads shorter than 50 bp. Trimmed paired reads were mapped to the *V. cholerae* N16961 reference genome (Accession number: chromosome 1 CP028827.1; chromosome 2 CP028828.1) using BWA-MEM v0.7.17 ^23^. Read count data was obtained with featureCounts from the Rsubread v2.10.2 package^24^. The expression levels, given in TPM values (transcripts per kilobase million) for all protein-coding genes, were calculated with the Geneious Prime “Calculate Expression Level” function and in R v4.2.2 (www.R-project.org) from FPKM values obtained with DESeq2 v1.36.0 ^25^. RNA-seq data are accessible through GEO Series accession number GSE229776 (https://www.ncbi.nlm.nih.gov/geo/query/acc.cgi?acc=GSE229776).

### Quantitative real-time PCR

Total RNA was extracted, as previously described, from three biological replicates of strains B118, B012, B014, B079, B016, B017, B084, and B088, grown in LB at 37°C for 20 h. To eliminate any residual DNA, the extracted RNA was treated using the TURBO DNA-free™ Kit (Invitrogen). RNA concentration was measured using a BioSpectrometer (Eppendorf). Prior to cDNA synthesis, a DNA wipe-out step was conducted using the QuantiTect® Reverse Transcription Kit (QIAGEN), following the manufacturer’s instructions. Subsequently, cDNA synthesis was carried out using the same kit with the following temperature steps: 42°C for 15 min, followed by 95 °C for 3 min. The obtained cDNA was diluted 10-fold and served as template in the qPCR.

RT-qPCR was performed in an Applied Biosystems QuantStudio 3 using the QuantiTect® Multiplex PCR Kit (QIAGEN). The primers and probes used are listed in Table S2. The qPCR protocol consisted of a HotStarTaq DNA Polymerase activation step at 95°C for 15 min, followed by 40 cycles of denaturation at 94°C for 60 s, and annealing/extension at 60°C for 60 s. Relative changes in the expression of the *gfp* and *mCherry* genes in all the tested strains, relative to the B118 strain (harboring the pDProm empty plasmid), were determined according to the threshold cycle method (2−ΔΔCT). The results were normalized against the expression of the housekeeping gene *gyrA*, as described in ^26^. Mean values were calculated from three independent biological replicates, each with three technical replicates.

### 5’ Rapid Amplification of cDNA Ends

Two micrograms of previously isolated RNA from strains B079, B016, B017, B084, and B088 were used to identify transcription start sites using the 5’ RACE System for Rapid Amplification of cDNA Ends, version 2.0 (Invitrogen), in accordance to the manufacturer’s protocol. The sequences of the primers used are listed in Table S2. First-strand cDNA was synthesized from total RNA using a gene-specific primer (GSP1). The cDNA products were subsequently amplified by PCR using Dream Taq DNA polymerase (Thermo Scientific), a nested gene-specific primer (GSP2) design to anneal to a site within the cDNA molecule, and a novel kit-supplied deoxyinosine-containing anchor primer. A cDNA control PCR was performed using sense gene-specific primer (GSP3), in conjunction with the GSP2 primer. If required, a second nested gene-specific primer (GSP4) was used for PCR amplification. PCR products were then analysed by agarose gel electrophoresis and purified using the GeneJET PCR purification kit (Thermo Scientific). The purified PCR products were subsequently cloned using the TOPO TA Cloning® kit (Invitrogen) and transformed into DH5α subcloning efficiency competent cells (Thermo Scientific), following the manufacturer’s instructions. At least six independent clones from each cassette were selected for DNA sequencing. The transcription start sites (+1 site) were determined by aligning the obtained sequences to the corresponding reference genome, being the +1 the nucleotide immediately following the poly(dC) tail.

### Bicyclomycin assay

To inhibit the activity of the Rho termination factor, a bicyclomycin assay was performed. Strains B118, B012, B014, B017, B079, B088, B084, and B016 were cultured on LB supplemented with kanamycin (75 µg/mL) for 20 h at 37°C. Samples were diluted 1:100 and bicyclomycin was added to the cultures at a concentration of 50 µg/mL at early exponential phase. After 2h-incubation at 37°C, cultures were diluted 1:400 on filtered NaCl (0,9 M) in 96-well plates (Nunc, Thermo Scientific) and analysed with a CytoFLEX-S flow cytometer (Beckman Coulter), as above described.

### Construction of the pDProm-*dfrB9* plasmid and derivatives

In order to test the inducibility of antibiotic resistance due to the presence of promoter-containing gene-less cassettes, the resistance cassette *dfrB9*, which confers resistance to trimethoprim, was selected from the IntegrALL database ^27^ and synthesized by IDT. To obtain the pDProm-*dfrB9* plasmid, the pDProm backbone was amplified excluding the *gfp* gene, while the *dfrB9* was amplified with 20 bp homology with the backbone. The *dfrB9* gene was then cloned into the backbone by Gibson assembly. The same procedure was performed with pDProm-Plac-*attC*-*gfp* (the *attC* region is not included in this plasmid version), and with pDProm-C7, -C8, -F3, -P4 and -V13 plasmids. The backbones were amplified excluding the *gfp* gene, and the *dfrB9* gene cassette was amplified with 20 bp homology with the corresponding backbone for further cloning downstream the gene-less fragments using Gibson assembly. The pDProm-*dfrB9* plasmid and all the derivatives were transformed in *V. cholerae* N16961 and *E. coli* DH5α following the above-described protocols.

For the cloning of cassettes F1 and C7 downstream the *dfrB9* in pDProm-C8-*dfrB9*, both cassettes F1 and C7 were amplified with 20 bp homology with the plasmid backbone. The plasmid pDProm-C8-*dfrB9* was amplified and cassettes F1 or C7 were then cloned into the backbone by Gibson assembly. The obtained plasmids pDProm-C8-*dfrB9*-F1 and pDProm-C8-*dfrB9*-C7 were transformed in *V. cholerae* N16961. Plasmids presence was confirmed by PCR with primers MRVII and MFD and verified by DNA sequencing. Primers used for pDProm-*dfrB9* plasmids construction are listed in Supplementary Table S2.

### Determination of the minimum inhibitory concentration (MIC) of trimethoprim

The MICs to trimethoprim of *V. cholerae* N16961 and *E. coli* DH5α containing pDProm-*dfrB9* and derivatives were performed following CLSI broth microdilution recommendations. Briefly, 10^5^ UFCs from an overnight culture were inoculated in 96-well plates (SPL Life Sciences) containing 200 µL of fresh MH (Oxoid) with doubling concentrations of trimethoprim (Sigma Aldrich). Plates were incubated at 37°C for 20 h without shaking. The MIC value was defined as the minimal concentration at which bacterial growth was inhibited in two biological replicates with two technical replicates each.

## RESULTS

### Identification of gene-less cassettes from *Vibrionaceae* genomes

We explored the data bases and selected four different genomes belonging to four different *Vibrio* species: *V. cholerae* N16961 (NZ_CP028828.1), *V. fischeri* ES114 (NC_006841), *V. parahaemolyticus* VPD14 (NZ_CP031781), and *V. vulnificus* FDAARGOS_663 (NZ_CP044069) with the aim of finding gene-less cassettes. The selection of these genomes was driven by several key factors: i) they contain SCIs; ii) these SCIs are of considerable length, with most of the array presumed to be silent if only the PC is active; and iii) experimental evidence exists for cassette recruitment by MIs ^28^.

We selected all cassettes lacking annotated genes, further focusing on those not encoding ORFs exceeding 210 bp (70 amino acid-long proteins) (see discussion section). According to these constrains, a total of 29 cassettes within the four selected species did not contain protein encoding genes. The distribution of these gene-less cassettes was as follows: nine in *V. cholerae* N16961; three in *V. fischeri* ES114; four in *V. parahaemolyticus* VPD14; and thirteen in *V. vulnificus* FDAARGOS_663. Additionally, we incorporated a cassette from *V. cholerae* N16961 that, although not containing an annotated gene, contained an ORF of a size slightly above the limit (78-amino acid). Altogether, our analysis yielded 30 cassettes.

### Construction and validation of the reporter plasmid pDProm

We designed a bidirectional reporter <u>p</u>lasmid for the <u>D</u>etection of <u>Prom</u>oter activity (pDProm) to use it with the gene-less cassettes from SCIs of *Vibrio* species. This plasmid contains two divergent promoterless genes encoding the fluorescent proteins GFP and mCherry, allowing for the simultaneous detection of promoter activity in both DNA strands of a given insert (Figure 2A). To validate the utility of pDProm, we wanted to mimic the structure of an integron cassette containing a promoter and verify its activity. To do so, we inserted a Plac promoter upstream a superintegron *attC* site -also known as *Vibrio cholerae* repeats (VCRs) ^29^- (the 128 bp *attC* site of cassette VCA0398 in *V. cholerae* N16961), in both orientations into pDProm (Figure 2B). We then introduced both plasmids in *V. cholerae* N16961 and we compared fluorescence intensity of GFP and mCherry in these strains with the values of the empty pDProm. As shown in Figure 2C, the Plac promoter increased GFP fluorescence 295-fold in Plac-*attC*-GFP, and mCherry intensity 60-fold in Plac-*attC*-mCherry. The lower fluorescence of the mCherry reporter is likely due to maturation or stability differences between fluorescent proteins. Indeed, unlike GFP, mCherry fluorescence does not increase during bacterial log phase and varies likely in response to different growth rates ^30^. However, a 60-fold increase is significant (*p*-value <0,0006) and provides a good dynamic range to test promoter activity. No significant differences were observed between the GFP and mCherry fluorescence raw values given by *V. cholerae* N16961 and the empty-pDProm strain (Supplementary Figure S2). We hence conclude that the use of pDProm is suitable for the detection of promoters in DNA fragments.

**Figure 2.**
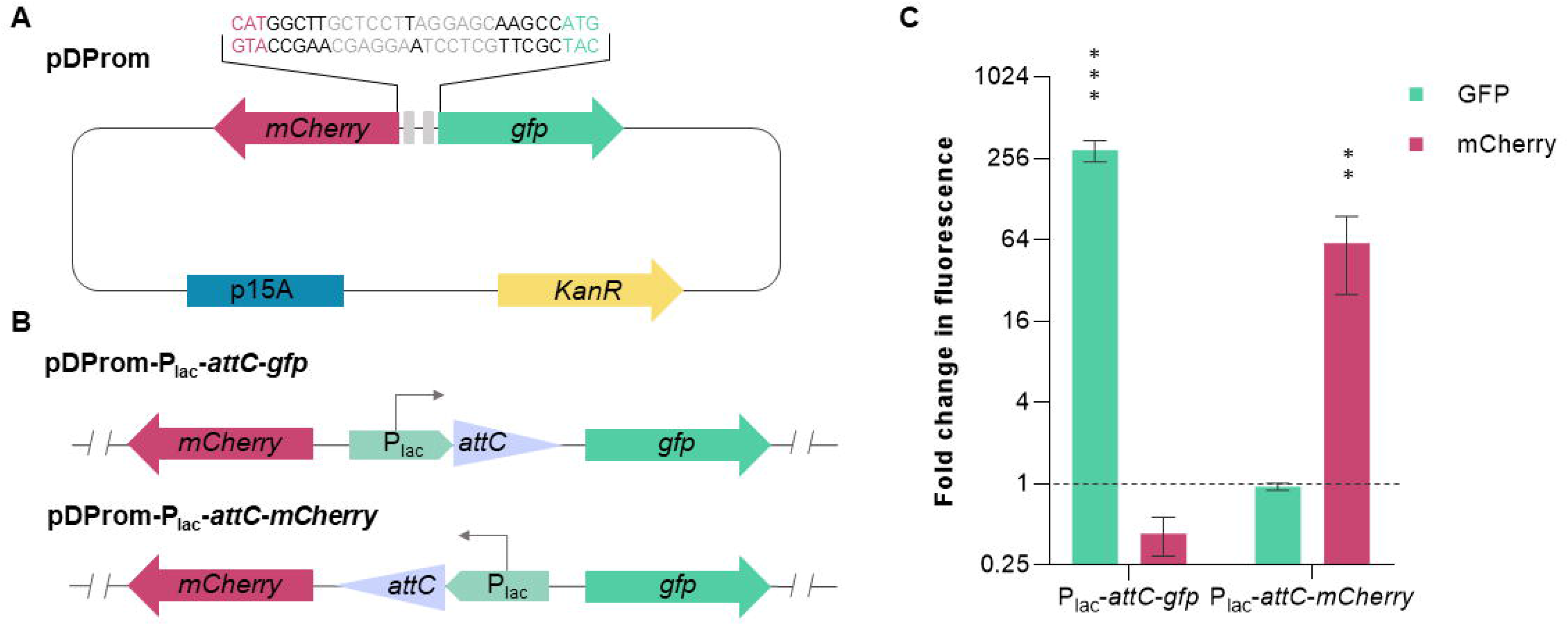
Structure and validation of the pDProm reporter plasmid. (**A**) Structure of the plasmid pDProm containing the reporter genes *mCherry* and *gfp*, the kanamycin resistance gene and the p15A origin of replication. The intergenic region sequence between *gfp* and *mCherry* is depicted including the ribosome binding sites (RBS) for both proteins (grey boxes and font) and the start codons (magenta and green font). (**B**) Schematic representation of the control DNA fragments. The pDProm-Plac-*attC*-*gfp* plasmid carries the *gfp* under the control of a Plac promoter upstream an *attC* site from *V. cholerae* N16961. The pDProm-Plac-*attC*-*mCherry* plasmid carries the *mCherry* under the control of a Plac promoter upstream an *attC* site from *V. cholerae* N16961. (**C**) Fold change values for fluorescence intensity given by pDProm-*attC*-*gfp* and pDProm-*attC*-*mCherry* relative to the the values obtained from the empty pDProm (dashed line). Error bars represent standard deviation of fluorescence measurements of three biological replicates with two technical replicates each. The *p*-values (*) were calculated by comparing each measure with that of the pDProm using paired t-test (** p 0,01-0001; *** p < 0,0001).

We synthesized the 30 gene-less cassettes identified previously with their respective *attC* sites and cloned them into pDProm. We introduced all plasmids in *V. cholerae* N16961 and verified the sequence of all constructs. For the sake of simplicity, cassette names are dubbed with the first letter of the species they belong to, and a correlative number.

### Determination of promoter activity

Using pDProm, cassettes containing a promoter on the sense strand will show an increase in GFP intensity, while those with a promoter on the antisense strand will present an increment in the mCherry fluorescence intensity. We have defined as *sense* strand the strand that would be under the influence of the PC promoter. From the ten cassettes selected from *V. cholerae* N16961, eight showed significant promoter activity on the sense strand, with 9 to 66-fold increases in GFP fluorescence. Among those with promoter activity, cassettes C5, C8, and C10 also showed significant promoter activity on the antisense strand, ranging from 2,8 to 3,5-fold (Figure 3A). Regarding the four gene-less cassettes from *V. parahaemolyticus* VPD14, three of them (P1, P2 and P4) showed 8 to 28-fold increases in GFP fluorescence. The P1 cassette also gave a 2,5-fold increase in the fluorescence intensity by the antisense strand (Figure 3B). After the screening of the thirteen gene-less cassettes from *V. vulnificus* FDAARGOS_663, we identified promoter activity on the sense strand in eleven of them (all but cassettes V7 and V12), with increment values varying from 5 to 20-fold in comparison with the empty pDProm. Besides, we also found significant promoter activity on the antisense strand in cassettes V2, V3, V11 and V12, all of them with around 2,5-fold increase in mCherry fluorescence intensity (Figure 3C). Finally, we analyzed the gene-less cassettes from the superintegron of *V. fischeri* ES114 and detected significant promoter activity in all of them on the sense strand with fold changes ranging from 27 to 101-fold. We also detected the highest promoter activity on the antisense strand in cassette F1 with a 14-fold increase in mCherry fluorescence (Figure 3D).

**Figure 3.**
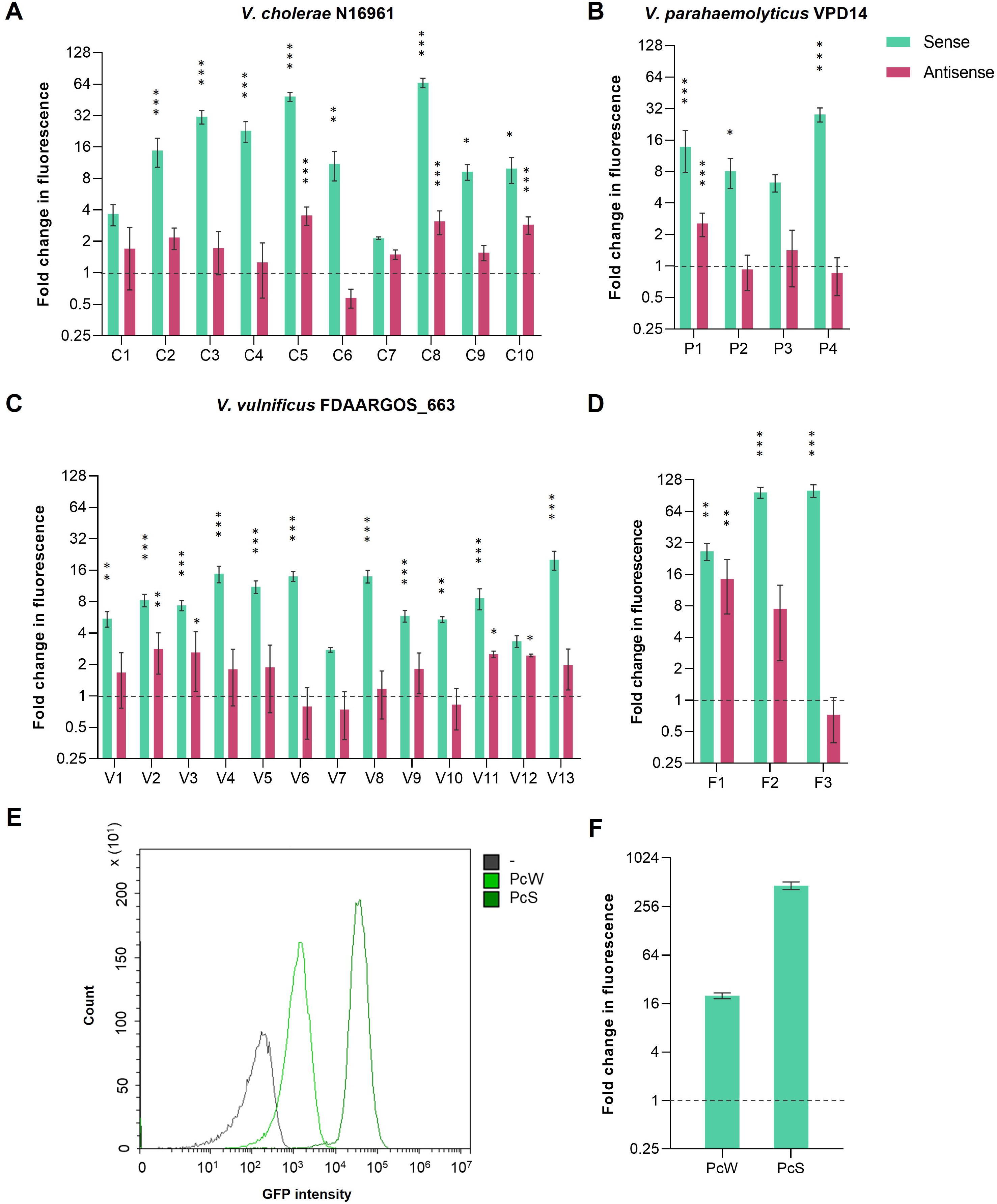
Determination of promoter activity of Vibrionaceae gene-less cassettes using pDProm. GFP (sense) and mCherry (antisense) fluorescence intensity fold change given by the selected cassettes from the superintegrons of (**A**) *V. cholerae* N16961, (**B**) *V. parahaemolyticus* VPD14, (**C**) *V. vulnificus* FDAARGOS_663, and (**D**) *V. fischeri* ES114, relative to the values given by the empty pDProm (dashed lines). (**E**) Fluorescence histograms and (**F**) fold change conferred by the PCW and PCS promoters in *V. cholerae* N16961 relative to the values given by the empty pDProm (dashed line). Error bars represent standard deviation of fluorescence measurements of three biological replicates with two technical replicates each. Histograms are the representation of one biological replicate. The p-values (*) were calculated by comparing each strain with the negative control strain B118 (empty pDProm) using one-way ANOVA with the Dunnett’s multiple comparisons test (* p 0,01-0,001; ** p 0,001-0,0001; *** p < 0,0001).

To validate our findings, we introduced three control DNA fragments from the *lacZ* gene into pDProm, with sizes corresponding to those of the tested gene-less cassettes (450-, 550-, and 650-bp), and measured the fluorescence intensity by flow cytometry. As expected, we observed no increase in the GFP or the mCherry expression levels when compared to the wildtype strain or the levels given by the empty pDProm vector (Supplementary Figure S3). According to these results, promoter activity seems to be prevalent among gene-less cassettes.

pDProm has a p15A origin of replication with 9 to 17 copies of the plasmid per cell ^31–33^. Hence, one could argue that the increases in fluorescence are due, at least in part, to the effect of copy number. So, in order to contextualize our results, we used a similar setup to measure the fluorescence levels of two variants of the PC promoter commonly found in Class 1 integrons: the PCS, which is considered a ‘strong’ promoter, and the PCW, a ‘weak’ promoter. Previous reports show that PCS is 30-fold stronger than PCW ^34,35^. We cloned these promoters upstream the *gfp* gene in the plasmid pA369 (also a p15A-based replicon), introduced them in *V. cholerae* N16961 and measured fluorescence by flow cytometry (Figure 3E). In our system, the fluorescence increments given by *V. cholerae* N16961 harboring the plasmids PcW-*gfp* or PcS-*gfp*, relative to the empty pDProm, were 19- and 525-fold, respectively (a 27-fold difference between both constructs, which is in accordance to the literature) (Figure 3F). By comparing these values with those obtained for the *Vibrionaceae* gene-less cassettes, we can conclude that most of them present a promoter strength similar to that of PCW, with the exception of cassettes C5, C8, F2 and F3, which have promoter strengths between PCW and PCS.

### Determination of promoter activity of gene-less cassettes by RT-qPCR

To demonstrate promoter activity at a transcriptional level, we performed RT-qPCR from a selected subset of strains, chosen for their diverse range of *gfp* and *mCherry* expression levels, previously measured by flow cytometry. We specifically targeted gene-less cassettes that exhibited robust promoter activity on the sense (F2, C5, and F1) and antisense strand (F1 and F2), alongside cassettes with mild or almost no such activity on both strands (V11 and V7, respectively) (Figure 4A and 4B). Transcription levels of both reporter genes were also measured in strains harboring pDProm-Plac-*attC*-*gfp* and pDProm-Plac-*attC*-*mCherry*, and were compared to the levels given by the empty vector. As shown in Figure 4C and 4D, we observed different transcription levels of the *gfp* and *mCherry* given by the expression of the selected cassettes. In most cases, the obtained transcription levels correlate well to the expression observed by measuring fluorescence, corroborating the consistency of our previous observations.

**Figure 4.**
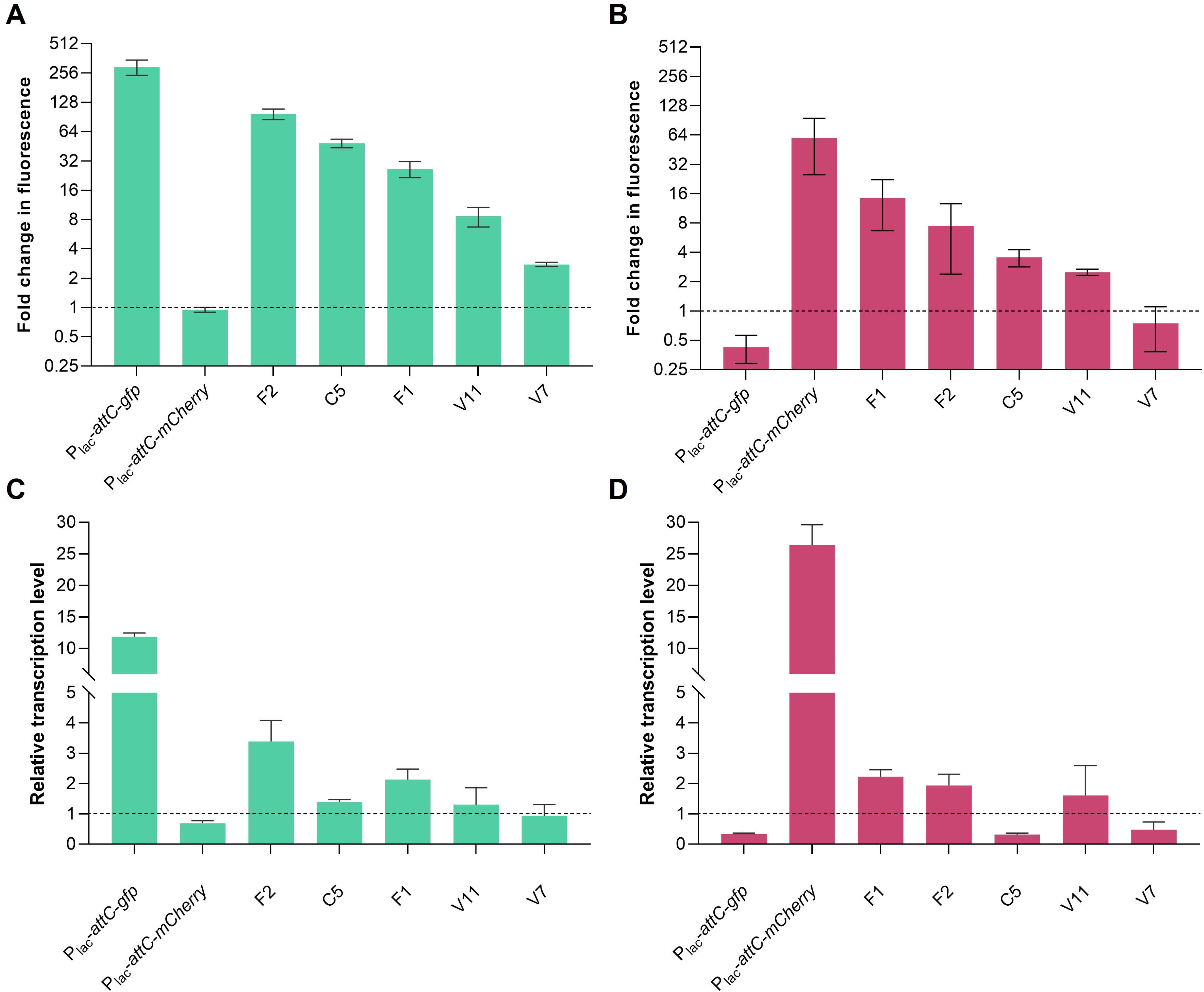
Validation of gene-less cassettes expression levels by RT-qPCR. (**A**) Fold change in fluorescence of GFP (sense) and (**B**) mCherry (antisense) given by the Vibrionaceae-selected cassettes relative to values obtained by the empty pDProm (dashed line). Error bars represent standard deviation of fluorescence measurements of three biological replicates with two technical replicates each. (**C**) Transcription levels of *gfp* and (**D**) *mCherry* determined by RT-qPCR from the selected Vibrionaceae gene-less cassettes relative to the empty pDProm. Error bars represent standard error of three independent biological replicates with three technical replicates each.

### Identification of transcription start sites in gene-less cassettes

Having confirmed promoter activity in specific *V. cholerae* gene-less cassettes by measuring the expression of the reporter genes *gfp* and *mCherry* by both flow cytometry and RT-qPCR, we proceeded to further characterize the potential transcription start sites (TSSs) (+1). To accomplish this, we performed the 5’ rapid amplification of cDNA ends (5’ RACE) technique, selecting the same gene-less cassettes (F1, F2, C5, V7, and V11) for which transcription activity had previously been validated via RT-qPCR. Following 5’ RACE, the resulting cDNA products were amplified with cassette-specific oligos and cloned into a pTOPO plasmid. Sequencing of a variety of clones revealed the TSSs positions for each selected gene-less cassette (Figure 5). Notably, TSSs were identified at different positions within the length of the cassette, and even within the *attC* recombination site, as in cassette F2. While certain cassettes exhibited a concentration of TSSs within a specific region, potentially indicating a single promoter (cassettes F1, C5, or V7), others displayed TSSs distributed at different points along the cassette at equal frequencies, as cassettes F2 and V11, suggesting a potential spurious transcription effect. Of particular interest, in a previous study Krin *et al*. located TSSs in six of the cassettes for which we have found promoter activity (C2, C3, C5, C6, C9 and C10). Our results for C5 are in accordance with the TSS at position 359 in cassette C5 ^36^, both validating our results and making reasonable to assume the rest of TSSs in other cassettes. Furthermore, we also performed the 5’ RACE assay on the antisense strand of cassette F1, where high expression levels were observed by measuring mCherry expression. As depicted in Figure 5, two main TSSs on the F1 antisense strand were identified, corroborating the presence of transcription on both strands in this gene-less cassette by distinct promoters.

**Figure 5.**
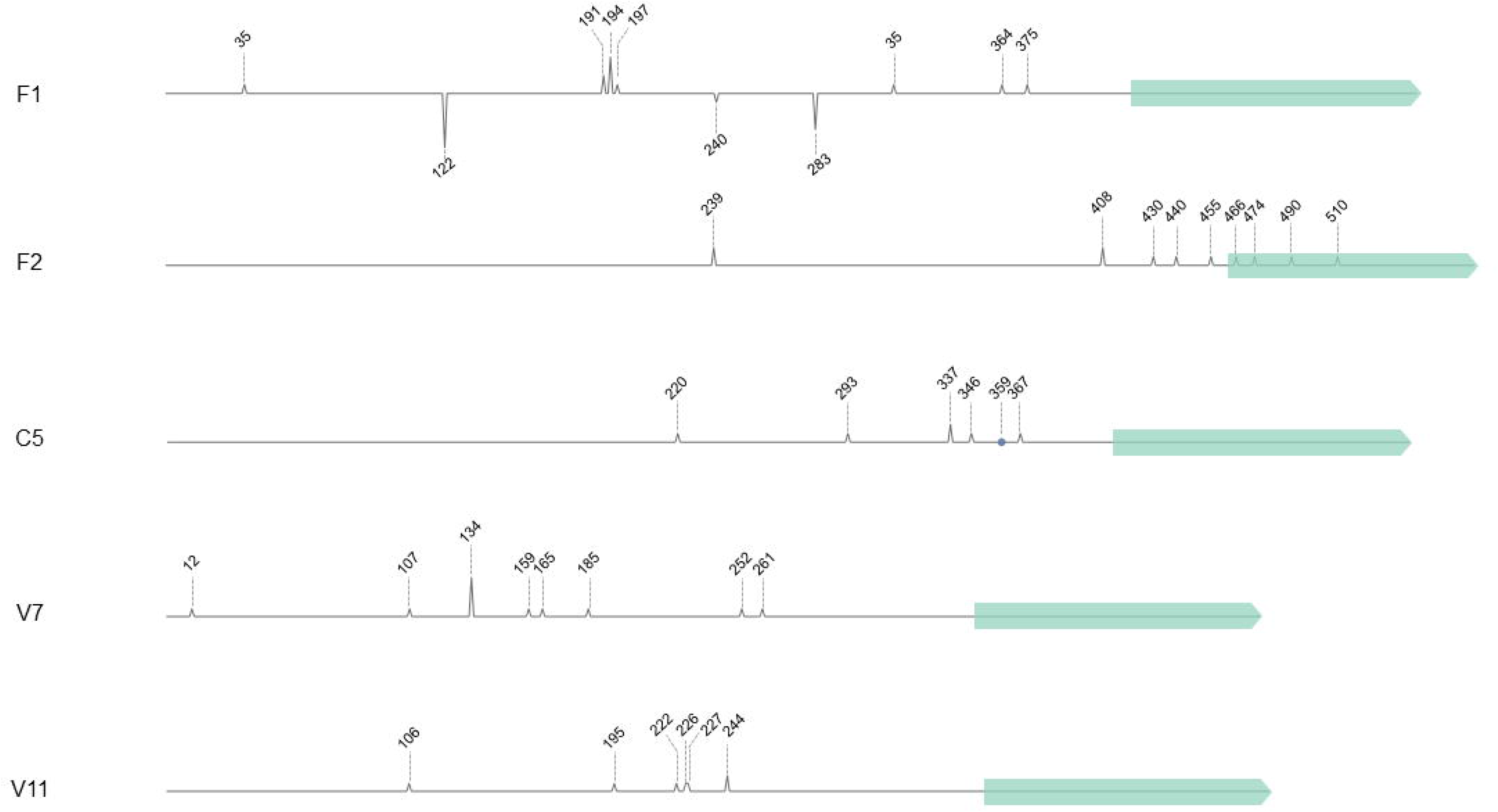
Transcription start sites (TSS) of Vibrionaceae gene-less cassettes. Schematic illustration of the gene-less cassettes F1, F2, C5, V7, and V11 from Vibrionaceae, indicating the detected TSSs identified through 5’ RACE. The height of the peaks corresponds to the frequency of detection for each TSS along the cassette. The peaks of cassette F1 below the line indicate the identified TSSs on the antisense strand. The blue point in cassette C5 represents a previously reported TSS position ^36^. Light green arrows represent the *attC* sites within each cassette.

Next, we wanted to understand whether spurious transcription is at play, and the potential role of the transcriptional terminator Rho -a terminator of spurious transcription ^37^. We measured gene expression in an assay using the Rho inhibitor bicyclomycin, and measured fluorescence levels by flow cytometry after 2h-incubation. Thus, if Rho is involved in the transcriptional termination in gene-less cassettes, its inhibition would result in increased fluorescence levels due to higher number of transcripts reaching the fluorescence reporter gene. Despite the inhibition of Rho, we did not observe any substantial enhancement in the GFP or mCherry fluorescence signals in comparison with the non-treated strains (Supplementary Figure S4), suggesting that Rho’s role in regulating cassette expression, if any, is absent or very limited in our conditions.

### Identification of promoters in gene-less cassettes cannot be performed *in silico*

Given the number of gene-less cassettes with promoter activity, we wanted to know whether this could have been accurately predicted *in silico*. To do this, we used BPROM software (Softberry.com) ^38^ and De Novo DNA ^39^ to predict the presence of putative σ^70^ promoters or transcription initiation rates, respectively, on both sense and antisense strands of the cassettes. BPROM identified promoters in both strands of all fragments tested and, in some cases, more than one promoter was predicted. The LDF (Linear Discriminant Function) score provided by BPROM is an indicator of promoter strength. The predicted -10 and -35 boxes and scores for all the cassettes are described in Supplementary Table S3. As controls, we also analyzed the previously described known promoters PCS, Plac and PCW, which had LDF values of 5.97, 3.94 and 3.10, respectively. We plotted, for all the cassettes, the LDF values for the sense (Supplementary Figure S5A) and antisense (Supplementary Figure S5B) strands along with the mean of GFP and mCherry fluorescence levels, respectively. We calculated the squared Pearson correlation coefficient and could not find a strong correlation between *in silico* and *in vivo* promoter strength (insets; sense strand: R^2^ = 0,037; *p* = 0,3; antisense strand R^2^ = 0,2; *p* = 0,01). The use of De Novo DNA software identified TSSs *in silico* that did not match those detected experimentally by 5’ RACE (Supplementary Figure S6). Generally, none of these algorithms would have allowed to identify *bona fide* promoters, confirming the need for experimental assessment.

### Transcription levels of integron cassettes from *V. cholerae* N16961 superintegron

Our results show that chromosomal integrons in *Vibrio* spp. have a wealth of gene-less cassettes with promoter activity scattered along the array. For instance, *V. cholerae* N16961 contains nine such cassettes, eight of which show at least half of the strength of a PCW and are therefore biologically relevant. These have to be added to the list of cassettes that encode a gene with a dedicated promoter, such as toxin-antitoxin systems, among others. It is therefore plausible that the operon-like model of expression is more complex on SCI, with many promoters along the array.

In order to investigate the levels of expression within the *V. cholerae* superintegron, and to contextualize the effect of gene-less cassettes, we performed RNA-seq for the *V. cholerae* strain N16961 in exponential and stationary growth phase conditions. We analyzed the data as described in Materials and Methods and expression levels of superintegron cassettes were measured calculating the number of transcripts per kilobase million (TPM), as depicted in Figure 6. We excluded from the analysis duplicated cassettes or those containing repeats of at least 300 bp, to avoid biased reporting of expression values of regions for which discrimination power is low. To provide an idea of physiological levels of gene expression, we also calculated the TPM values for 8 housekeeping genes. As a validation of our results, the sigma factor RpoS shows higher levels of expression in stationary phase, as expected.

**Figure 6.**
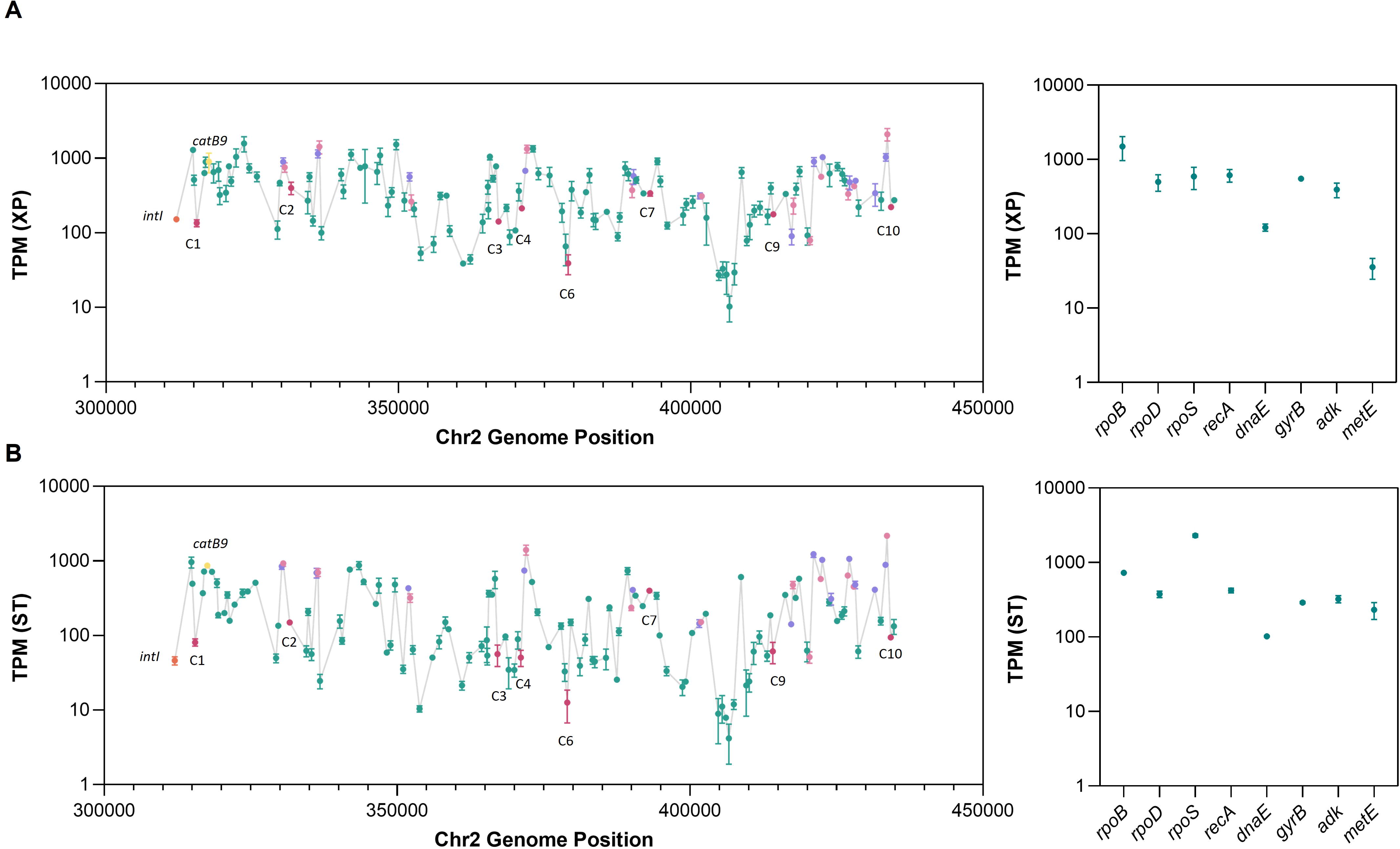
Expression levels of *V. cholerae* N16961 superintegron gene cassettes. (**A**) Expression levels represented as transcripts per kilobase million (TPM) of gene cassettes along the superintegron array, and eight housekeeping genes used as a reference, were calculated from RNA-seq data obtained from exponential (XP) and (**B**) stationary (ST) growth conditions. TPM values of genes-less cassettes (magenta), toxin-antitoxin systems (mauve and pink), integrase (orange), and the chloramphenicol resistance gene catB9 (yellow), are highlighted. Error bars represent the standard deviation of three independent biological replicates.

Our data show that -in addition to TA systems that contain their own promoter-, many cassettes are expressed at detectable levels. TPM values show broad variation along the array, with 210-fold and 500-fold differences between the most and least expressed cassettes in exponential (Figure 6A), and stationary phase, respectively (Figure 6B). Regions or “blocks” of gene cassettes that present similar expression levels are also detected throughout the length of the superintegron. The expression levels of many of the cassettes are in the same range as those of housekeeping genes, proving the physiological relevance of our measures. Our results were further verified by analyzing the data from an independent RNAseq from the same strain available in the databases (BioProject PRJNA420494) ^22^. We found a similar distribution of reads along the array (Supplementary Figure S7), allowing to reach the same general conclusions.

To reveal the role of gene-less cassettes in their natural context we took a closer look at the mapping of transcripts of the 6 cassettes with expression levels similar to PcW (C2 to C6 and C8). Our analysis found technical and biological limitations. Cassettes C5 and C8 contained repeated regions and were discarded to avoid mapping biases; and cassette C3 could not be clearly interpreted (Figure 7B) because it is located immediately upstream of the parD/E3 toxin-antitoxin system encoding a promoter in the opposite orientation. Of the remaining cassettes, C4 and C6 showed only mild increases in transcript levels (Supplementary Figure S8), while a clear increase in transcription could be observed inside the sequence of C2 (Figure 7A), supporting its role as promoter of downstream cassettes. Our results indicate that the superintegron is not a “silent” structure and that the expression of gene cassettes in the array does not rely solely on the PC promoter. We also find evidence of a gene-less cassette acting as a promoter in its native context.

**Figure 7.**
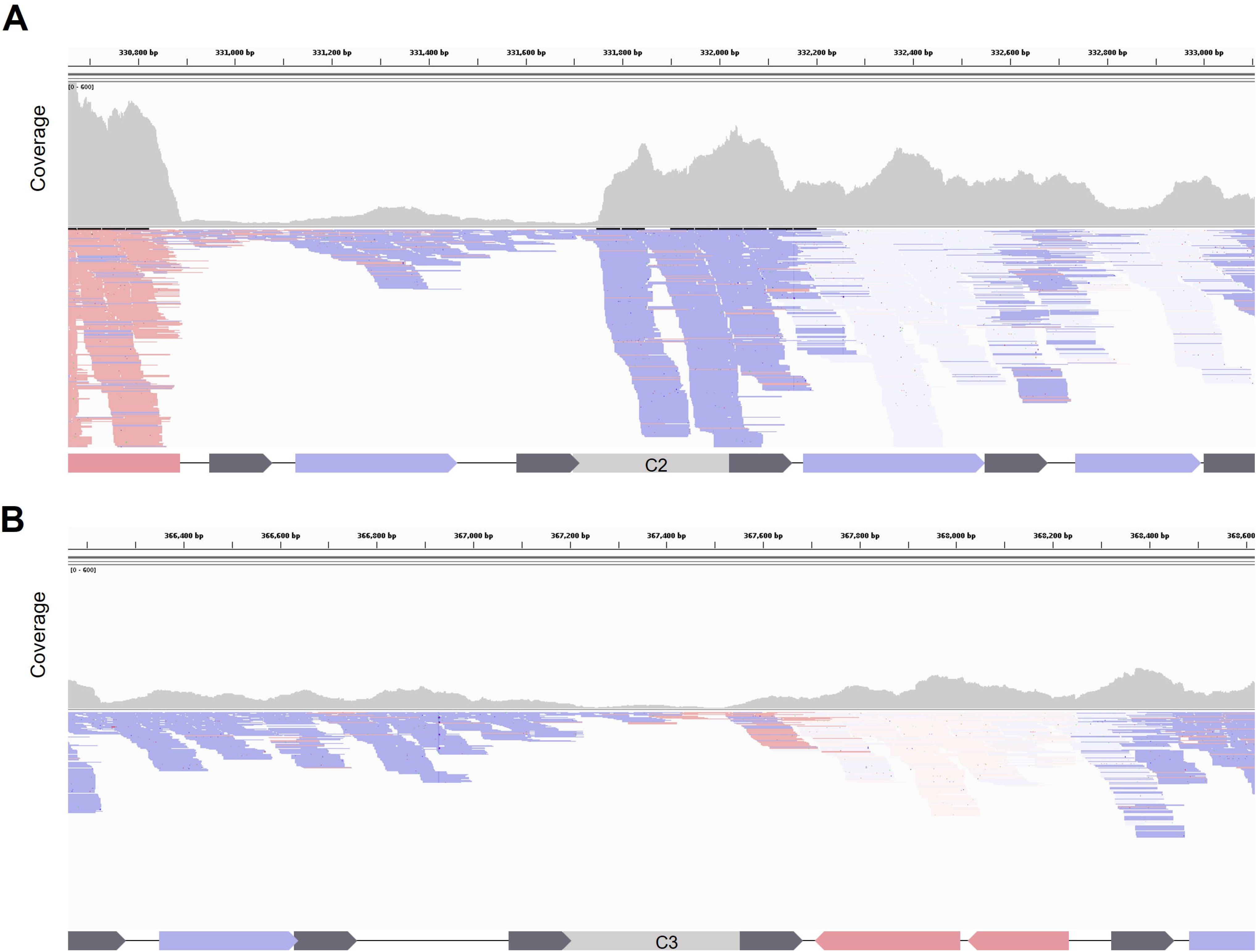
Representation of reads from the RNAseq of *V. cholerae* N16961 superintegron. (**A**) The number and orientation of reads, and the coverage upstream and downstream gene-less cassettes C2 and (**B**) C3 were analyzed by Integrative Genomics Viewer (IGV) software. Blue color designates genes and reads from the plus (sense) strand; red color designates genes and reads from the minus (antisense) strand; uncolored reads indicate unknown status or reads repetitions.

### Promoter-carrying gene-less cassettes on the sense strand can trigger antibiotic resistance

Given the possibility that mobile integrons recruit cassettes from SCIs with promoter activity, we aimed to investigate their potential role in promoting antibiotic resistance when located upstream of a silent resistance gene. To test this, we selected the plasmids containing the gene-less cassettes with the highest promoter activity from each *Vibrio* species (C8, P4, V13, and F3) and replaced the *gfp* gene with the promoterless trimethoprim resistance cassette *dfrB9*. As negative and positive controls, we constructed pDProm-*dfrB9* and pDProm-Plac-*dfrB9*, respectively (Figure 8A). Last, we also tested the C7 cassette from *V. cholerae* N16961, that encodes a putative 78-amino acid long protein. As we did not detect fluorescence increase in either the C7 sense or antisense strands (Figure 3A), we used this promoterless cassette as a control. We introduced all plasmids in *V. cholerae* N16961 and performed MIC assays to trimethoprim. We found that the four selected gene-less cassettes increased trimethoprim resistance from two- and four-fold (cassettes C8 and V13) to more than 500-fold in the case of P4 and F3 (Figure 8B). The same result was obtained for the positive control Plac-*dfrB9,* while strains with C7 or the empty pDProm-*dfrB9* showed the same level of resistance as the strain with no plasmid.

**Figure 8.**
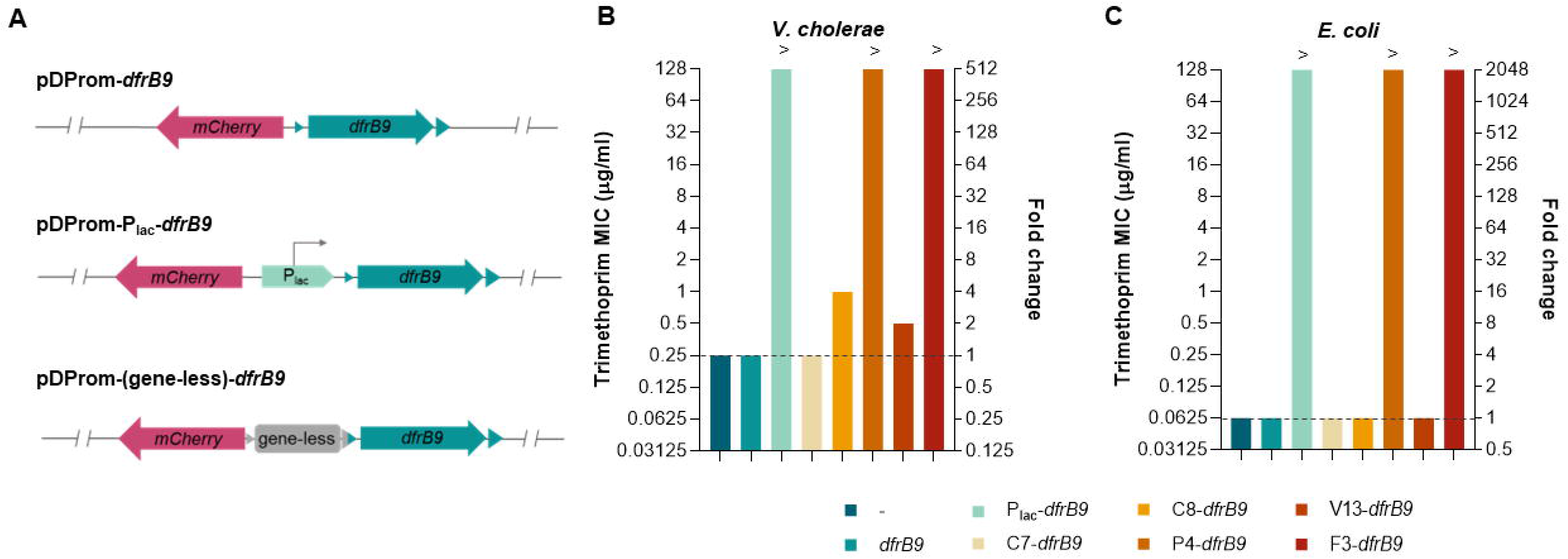
Promoter-carrying gene-less cassettes induce trimethoprim resistance. (**A**) Fragments of the pDProm-*dfrB9* plasmids where the *gfp* gene is replaced by the trimethoprim resistance cassette *dfrB9* in pDProm-Plac and pDProm-“gene-less” plasmids. (**B**) Minimum inhibitory concentration (MIC) to trimethoprim and resistance fold change of *V. cholerae* N16961 and (**C**) *E. coli* DH5α containing pDProm-*dfrB9* plasmids with the cassettes yielding the highest fluorescence intensity from each *Vibrio* (C8, P4, V13 and F3). Strains containing Plac-*dfrB9* or pDProm-*dfrB9* (*dfrB9*, empty vector) were used positive and negative controls of trimethoprim resistance, respectively. C7-*dfrB9* was used as promoterless control. The resistance levels of *V. cholerae* N16961 and *E. coli* DH5α strains with no plasmids (-) are indicated with a dashed line. MIC values above 128 μg/ml are represented with (>). The MIC of each strain was averaged from two biological replicates with two technical replicates each.

Since ICs can be mobilised and transferred through horizontal gene transfer among unrelated species, we were interested in the effect that these gene-less cassettes may have in the resistance levels when present in a different species. To address this, we transformed these plasmids in *E. coli* and measured the resistance levels to trimethoprim. In accordance to what we observed in *V. cholerae,* cassettes P4, F3 and the Plac-*dfrB9* control showed 2,000-fold increases in trimethoprim resistance, while strains with the C7 control, or the empty pDProm-*dfrB9* plasmid, showed no differences with the empty strain. These results suggest that promoter-carrying gene-less cassettes, when located upstream a resistance gene cassette in a plasmid, are able to induce high levels of resistance not only in vibrios, but also in more phylogenetically distant species as *E. coli*.

### Promoter-carrying gene-less cassette on the antisense strand reduce antibiotic resistance levels

Given that promoter activity was also detected on the antisense strand of some cassettes, we decided to test their influence on the expression of upstream cassettes. We hypothesize that if antisense promoters are functional, we would observe a decrease in trimethoprim resistance if cloned downstream *dfrB9*. We selected cassette F1 from *V. fischeri*, which showed a 14-fold increase in mCherry fluorescence and where two TSSs have been detected, and cloned it in pDProm-C8-*dfrB9* downstream *dfrB9*. It is of note that in this setup *dfrB9* expression is dependent on the C8 cassette. As a control, we also cloned the cassette C7 downstream *dfrB9* (Figure 9A), which did not show promoter activity. We introduced both plasmids in *V. cholerae* and measured the MIC to trimethoprim. After 24 h (Figure 9B), we observed that the MICs of strains where *dfrB9* was expressed (pDProm-C8-*dfrB9* and pDProm-C8-*dfrB9*-C7) were 2-to 4-fold higher than those of strains without plasmid or carrying a silent *dfrB9*. In the presence of F1, the MIC decreased to the same levels as in the case where *dfrB9* is silent or absent (Figure 9B). This effect was accentuated after 48 h of incubation, when strains expressing *dfrB9* reached above 8-fold higher MICs, while controls and the strain carrying the pDProm-C8-*dfrB9*-F1 retained the same values (Figure 9C). To better characterize these differences, we also performed growth curves of *V. cholerae* with all plasmids in the presence of inhibitory concentrations of trimethoprim (0,25 and 0,5 μg/ml). As observed in Figure 9D, all strains grew similarly in MH, but only those carrying pDProm-C8-*dfrB9* or pDProm-C8-*dfrB9*-C7 were able to grow in the presence of the antibiotic. Only after 32 h of incubation, one out of four replicates of *V. cholerae* with pDProm-C8-*dfrB9*-F1 started growing, probably due to the presence of spontaneous mutations. These data show, for the first time, that the presence of a promoter on the antisense strand of a gene-less cassette can suppress the expression of a gene, leading, in this case, to a reduction of antibiotic resistance.

**Figure 9.**
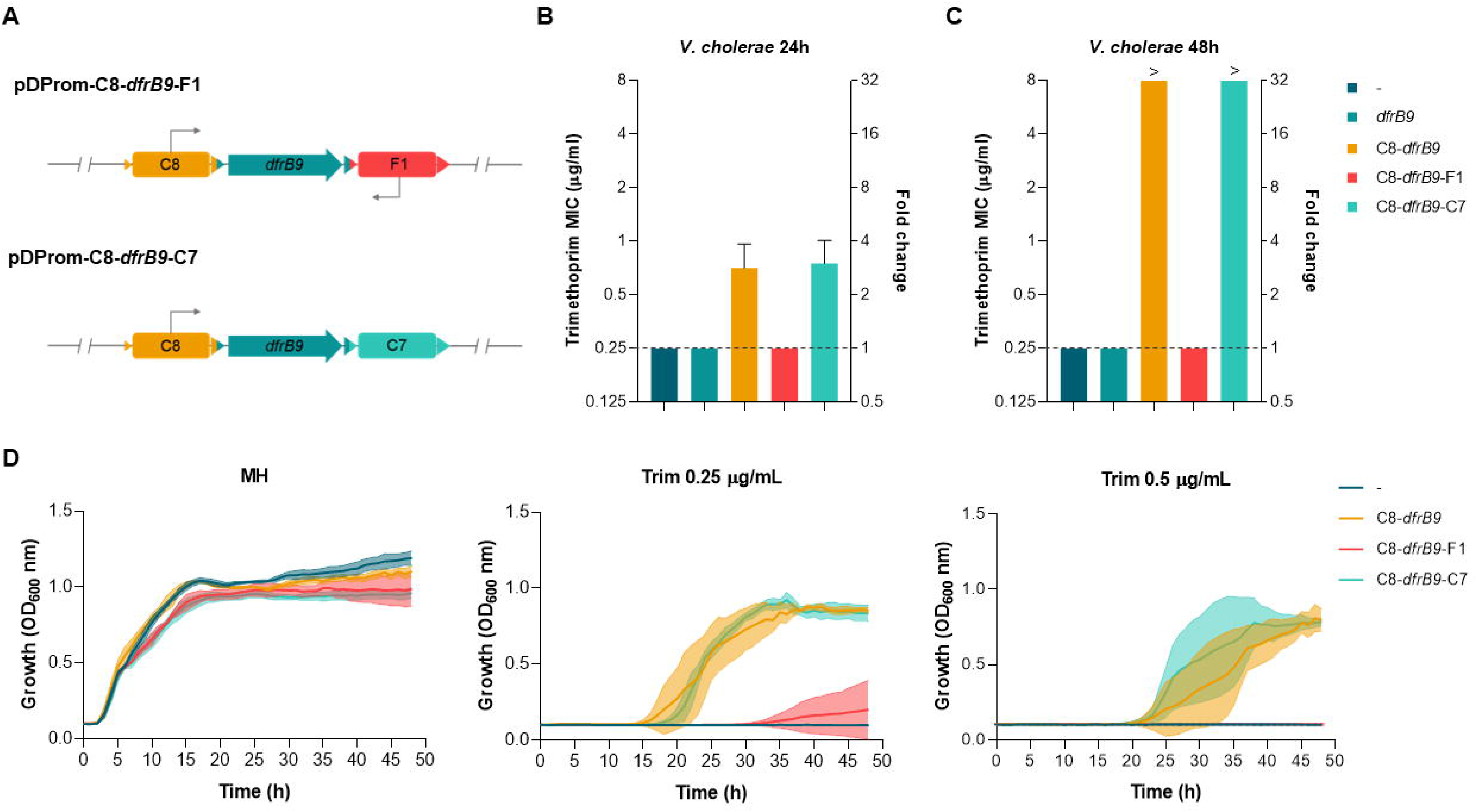
Gene-less cassette carrying a promoter on the antisense strand act as a transcriptional terminator. (**A**) Fragments of the plasmid pDProm-C8-*dfrB9* where cassette F1, as a cassette containing a promoter on the antisense strand, and C7, as a promoterless cassette control, are cloned downstream *dfrB9* (pDProm-C8-*dfrB9*-F1 and pDProm-C8-*dfrB9*-C7). (**B**) Minimum inhibitory concentration (MIC) to trimethoprim and resistance fold change at 24 h and (**C**) 48 h of *V. cholerae* N16961 containing plasmids pDProm-C8-*dfrB9*, pDProm-C8-*dfrB9*-F1 or pDProm-C8-*dfrB9*-C7. The strain containing pDProm-*dfrB9* (*dfrB9*, empty vector) was used a negative controls of trimethoprim resistance. The resistance levels of *V. cholerae* N16961 with no plasmids (-) are indicated with a dashed line. MIC values above 8 μg/ml are represented with (>). (**D**) Growth curves of *V. cholerae* N16961 containing plasmids pDProm-C8-*dfrB9*, pDProm-C8-*dfrB9*-F1 or pDProm-C8-*dfrB9*-C7 in the absence (MH) or presence of trimethoprim at 0,25 and 0,5 μg/ml. Error bars represent standard deviation of at least three independent biological replicates.

## DISCUSSION

The *Vibrionaceae* family comprises bacteria that are ubiquitously distributed throughout aquatic environments and includes important pathogens. *V. cholerae* is the causative agent of Cholera disease, that still causes approximately 100,000 deaths worldwide annually ^40^; while *V. parahaemolyticus* and *V. vulnificus* cause acute gastroenteritis, wound necrosis and/or sepsis in humans due to contaminated water or seafood consumption ^41,42^. A notable characteristic of the *Vibrionaceae* family is the presence of large SCIs in their genomes, carrying a highly diverse repertoire of ICs ^43,44^. These have likely been important agents in the evolution of vibrios, a phenomenon for which knowledge is scarce ^45^. It is therefore important to decipher the functions of ICs by revealing the role of the proteins they encode, yet, previous studies have shown that a considerable proportion do not appear to encode any (from 6% in *V. cholerae* to 49% in *V. vulnificus* CMCP6) ^46^. In this study, we have experimentally detected promoter activity in gene-less cassettes from the sedentary chromosomal integrons of four *Vibrio* species.

The general integron model posits that the PC promoter drives the expression of cassettes and that the increasing distance to the PC causes their silencing. Some exceptions to this rule are known, mainly in the form of cassettes encoding genes with dedicated promoters. The presence of promoters in cassettes within SCIs, and their impact on adjacent cassettes have seldom been investigated, although their presence has been previously suggested ^47^. In a study of the array of *V. rotiferianus*, the authors detected through RT-qPCR patterns of *en bloc* transcription along the array, suggesting the presence of internal promoters ^48^. In Krin *et al*.’s work, the authors performed the mapping of the transcription start sites (TSSs) in *V. cholerae* N16961, showing that many TSSs are scattered along the superintegron ^36^. In both studies, the origin of promoter activity was not linked to a specific type of cassette. Tansirichaiya *et al*. ^19^ were the firsts to report promoter activity in gene-less cassettes of *Treponema spp.* from the human oral metagenome ^49^. It is worth noting that this work only characterized 3 cassettes and that SCIs in *Treponema* are atypical because they encode the integrase in the same strand as the cassettes ^50,51^. Therefore, it is difficult to know if the presence of promoter cassettes is extrapolable to integrons in general. Here, we aimed to provide additional experimental evidence on the presence of such cassettes in canonical integrons. We have done so by tackling three limitations of the previous work. First, we test a larger number of cassettes (30 *vs*. 3) from a variety of species, providing a better grasp on the universality of the phenomenon. Second, in the previous study, the promoter activity of *Treponema spp.* cassettes was tested in *E. coli*, a very distantly related species. It is hence possible that the promoters identified might not be acting as such in the original host. Here, we test promoter activity in *V. cholerae,* which is the host of several cassettes, and it is a species closely related to the other hosts. Third and last, we confirm our fluorescence results through RNA-seq, RT-qPCR and 5’ RACE, and we explore the biological consequences of the presence of such promoters. We demonstrate their importance in modulating antimicrobial resistance provided by the *dfrB9* IC and prove that this effect is conserved in enterobacteria, where mobile integrons are commonplace and important actors in multidrug resistance. It is of note that we show for the first time that antisense promoters can interfere with the expression of upstream cassettes, acting as array silencers. These data also confirm the previous statement that *attC* sites do not act as transcriptional terminators ^52^. These results evidence the impact that such cassettes could have on AR if mobilized. Interestingly, some MIs contain <u>g</u>ene <u>c</u>assettes of <u>u</u>nknown function (*gcu*), many of which are very small and probably do not contain genes. Thus, it is possible that promoter cassettes are already modulating resistance genes in clinical settings, a matter that needs further investigation.

An important aspect of this and previous works is the definition of gene-less (or ORF-less) cassettes. Here, we have defined them as cassettes lacking annotated genes and not containing ORFs longer than 210 bp. A limitation of our work (and others) is that we have not experimentally proven that smaller ORFs do not encode proteins (nor small RNAs for that matter). However, there are several reasons why we believe our definition is sensible. First, cassette architecture is generally very compact, with ORFs occupying most -if not all-of the cassette sequence, sometimes even overlapping the *attC* site. The cassettes selected here range in size from 491 to 666 bp and *attC* sites of 101 to 127 bp. This leaves coding regions of sizes between 335 and 515 bp. Hence, the presence of protein coding genes smaller than 210 bp would be at odds with the general compaction of cassette coding. Second, we performed a search with BLASTP suite using the complete sequence of all cassettes that allowed us to confirm the absence of small peptides with high degrees of conservation. Surprisingly, in a few cases, we found that some gene-less cassettes contained truncated proteins that are still conserved intact in other genomes in the databases. The clearest example is cassette P3, that yields hits with 90% identity to a 128 aa-long protein from *V. alginolyticus*, but the sequence in P3 contains three premature stop codons in that frame. This BLASTP search supports our premise that cassettes selected here are gene-less and also suggests that -at least some of them-derive from gene-encoding cassettes that have degenerated. Additionally, the fact that TSSs in our results are not found at the 5’ end of cassettes suggests that these were not promoters dedicated to the expression of a gene in the cassette prior to pseudogenization. Altogether, this is surprising due to the fact that it is generally accepted that cassettes do not contain pseudogenes, and this is interpreted as proof of selective pressure acting upon them frequently enough to avoid pseudogenization ^53^. Because the loss of useless cassettes is especially easy in integrons (through excision reactions), it is possible that the function as promoters provides an adaptive value to these cassettes that saves them from purifying selection.

We have attempted to show the promoter activity of cassettes in their natural context by analyzing transcription rates of these cassettes in *V. cholerae*’s superintegron through RNA-seq. Despite technical and biological limitations, we could clearly observe the promoter activity of C2 (Figure 7A). Interestingly, the activity of C3 was possibly masked by the presence of the parD/E3 toxin-antitoxin cassette downstream, encoding a promoter in opposite orientation (Figure 7B). Thus, our RNA-seq data, and other datasets available, support that the array is not generally silent, with expression levels being very variable, and therefore, that there must be promoters along the array.

Our 5’-RACE results show that some cassettes contain a small subset of well-defined TSSs, while others exhibit multiple TSSs distributed throughout the cassette structure, including the *attC* site. The latter observation could be reflecting spurious transcription, possibly influenced by a high AT-content, rendering the DNA sequences similar to the promoters -10 elements (Supplementary Figure S6) ^54^. Cassettes are indeed known to be AT-rich compared to the rest of the genome, as is common among mobile genetic elements ^55^. Inhibition of Rho did not change the expression patterns, so the question on how transcription starts and is regulated in these cassettes remains uncertain. Nevertheless, comparison with mock DNA controls show that, irrespective of the mechanism, these findings are biologically relevant and have strong implications in the working model of long SCIs. Regarding cassette F1, with antisense promoter activity, we identified two specific locations where the majority of TSSs colocalize, but not a bidirectional promoter. The presence of promoter activity in both strands, puts forward the possibility of cassettes silencing the array upstream and adds a layer of complexity to the regulatory landscape of integrons ^56,57^.

In summary, our results indicate that gene-less cassettes in *Vibrionaceae* species encode promoters, extending previous observations on *Treponema* ICs to canonical integrons. This has several implications: first, it opens a new functional category in ICs -“promoter cassettes”-, expanding our understanding of cassette functions. Second, it confirms that several regions of SCIs are likely expressed in every array, enhancing the chances of finding adaptive cassettes in the array when needed, but likely increasing the cost of the structure ^48^. Third, these promoter cassettes can have a major impact in antibiotic resistance in different species if mobilized, a phenomenon that might have been neglected.

## DATA AVAILABILITY

The RNA-seq data discussed in this publication have been deposited in NCBI’s Gene Expression Omnibus ^58^ and are accessible through GEO Series accession number GSE229776 (https://www.ncbi.nlm.nih.gov/geo/query/acc.cgi?acc=GSE229776).

## SUPPLEMENTARY DATA

Supplementary Data are available at NAR online.

## AUTHOR CONTRIBUTIONS

Paula Blanco: Conceptualization, Formal analysis, Methodology, Validation, Writing-original draft. Alberto Hipólito: Methodology, Writing-review and editing. Lucía García-Pastor: Formal analysis, Methodology, Validation. Filipa Trigo: Methodology, Writing-review and editing. Laura Toribio-Celestino: Methodology, Validation, Writing-review and editing. Cristina Ortega: Methodology. Ester Vergara: Methodology. Álvaro San Millán: Conceptualization, Formal analysis, Writing-review and editing. José Antonio Escudero: Conceptualization, Formal analysis, Validation, Writing-original draft.

## REFERENCES

1. Mazel, D. Integrons: Agents of bacterial evolution. Nature Reviews Microbiology vol. 4 Preprint at 10.1038/nrmicro1462 (2006).

2. Boucher, Y., Labbate, M., Koenig, J. E. & Stokes, H. W. Integrons: mobilizable platforms that promote genetic diversity in bacteria. Trends Microbiol 15, (2007).

3. Gillings, M. R. Integrons: Past, Present, and Future. Microbiology and Molecular Biology Reviews 78, (2014).

4. Hipólito, A. et al. The expression of aminoglycoside resistance genes in integron cassettes is not controlled by riboswitches. Nucleic Acids Res 50, 8566–8579 (2022).

5. Cury, J., Jové, T., Touchon, M., Néron, B. & Rocha, E. P. Identification and analysis of integrons and cassette arrays in bacterial genomes. Nucleic Acids Res 44, (2016).

6. Zhu, Y.-G. et al. Microbial mass movements. Science (1979) 357, 1099–1100 (2017).

7. Rowe-Magnus, D. A. et al. The evolutionary history of chromosomal super-integrons provides an ancestry for multiresistant integrons. Proceedings of the National Academy of Sciences 98, 652–657 (2001).

8. Escudero*, J. A., Loot*, C., Nivina, A. & Mazel, D. The Integron: Adaptation On Demand. Microbiol Spectr 3, (2015).

9. Stokes, H. W., O’Gorman, D. B., Recchia, G. D., Parsekhian, M. & Hall, R. M. Structure and function of 59-base element recombination sites associated with mobile gene cassettes. Mol Microbiol 26, (1997).

10. Hall, R. M. & Stokes, H. W. Integrons: Novel DNA elements which capture genes by site-specific recombination. Genetica 90, (1993).

11. Collis, C. M. & Hall, R. M. Site-specific deletion and rearrangement of integron insert genes catalyzed by the integron DNA integrase. J Bacteriol 174, (1992).

12. Collis, C. M. & Hall, R. M. Gene cassettes from the insert region of integrons are excised as covalently closed circles. Mol Microbiol 6, (1992).

13. Collis, C. M. & Hall, R. M. Expression of antibiotic resistance genes in the integrated cassettes of integrons. Antimicrob Agents Chemother 39, (1995).

14. Biskri, L. & Mazel, D. Erythromycin esterase gene ere(A) is located in a functional gene cassette in an unusual class 2 integron. Antimicrob Agents Chemother 47, (2003).

15. Stokes, H. W. & Hall, R. M. Sequence analysis of the inducible chloramphenicol resistance determinant in the TN1696 integron suggests regulation by translational attenuation. Plasmid 26, (1991).

16. da Fonseca, É. L. & Vicente, A. C. P. Functional characterization of a cassette-specific promoter in the class 1 integron-associated qnrVC1 gene. Antimicrob Agents Chemother 56, (2012).

17. Holmes, A. J. et al. The gene cassette metagenome is a basic resource for bacterial genome evolution. Environ Microbiol 5, (2003).

18. Stokes, H. W. et al. Gene Cassette PCR: Sequence-Independent Recovery of Entire Genes from Environmental DNA. Appl Environ Microbiol 67, (2001).

19. Tansirichaiya, S., Mullany, P. & Roberts, A. P. Promoter activity of ORF-less gene cassettes isolated from the oral metagenome. Sci Rep 9, (2019).

20. Tansirichaiya, S., Rahman, M. A., Antepowicz, A., Mullany, P. & Roberts, A. P. Detection of Novel Integrons in the Metagenome of Human Saliva. PLoS One 11, e0157605 (2016).

21. Gibson, D. G. et al. Enzymatic assembly of DNA molecules up to several hundred kilobases. Nat Methods 6, (2009).

22. Sepúlveda-Cisternas, I., Lozano Aguirre, L., Fuentes Flores, A., Vásquez Solis de Ovando, I. & García-Angulo, V. A. Transcriptomics reveals a cross-modulatory effect between riboflavin and iron and outlines responses to riboflavin biosynthesis and uptake in Vibrio cholerae. Sci Rep 8, 3149 (2018).

23. Li, H. Aligning sequence reads, clone sequences and assembly contigs with BWA-MEM. (2013) doi:10.48550/arxiv.1303.3997.

24. Liao, Y., Smyth, G. K. & Shi, W. featureCounts: an efficient general purpose program for assigning sequence reads to genomic features. Bioinformatics 30, 923–930 (2014).

25. Love, M. I., Huber, W. & Anders, S. Moderated estimation of fold change and dispersion for RNA-seq data with DESeq2. Genome Biol 15, 550 (2014).

26. Liu, X., Beyhan, S., Lim, B., Linington, R. G. & Yildiz, F. H. Identification and Characterization of a Phosphodiesterase That Inversely Regulates Motility and Biofilm Formation in *Vibrio cholerae*. J Bacteriol 192, 4541–4552 (2010).

27. Moura, A. et al. INTEGRALL: a database and search engine for integrons, integrases and gene cassettes. Bioinformatics 25, 1096–1098 (2009).

28. Rowe-Magnus, D. A., Guerout, A.-M. & Mazel, D. Bacterial resistance evolution by recruitment of super-integron gene cassettes. Mol Microbiol 43, 1657–1669 (2002).

29. Mazel, D., Dychinco, B., Webb, V. A. & Davies, J. A Distinctive Class of Integron in the *Vibrio cholerae* Genome. Science (1979) 280, 605–608 (1998).

30. Cooper, K. G., Chong, A., Starr, T., Finn, C. E. & Steele-Mortimer, O. Predictable, tunable protein production in Salmonella for studying host-pathogen interactions. Front Cell Infect Microbiol 7, (2017).

31. Hipólito, A., García-Pastor, L., Vergara, E., Jové, T. & Escudero, J. A. Profile and resistance levels of 136 integron resistance genes. npj Antimicrobials and Resistance 1, 13 (2023).

32. Shao, B. et al. Single-cell measurement of plasmid copy number and promoter activity. Nat Commun 12, 1475 (2021).

33. San Millan, A., Escudero, J. A., Gifford, D. R., Mazel, D. & MacLean, R. C. Multicopy plasmids potentiate the evolution of antibiotic resistance in bacteria. Nat Ecol Evol 1, 0010 (2016).

34. Jové, T., Da Re, S., Denis, F., Mazel, D. & Ploy, M.-C. Inverse Correlation between Promoter Strength and Excision Activity in Class 1 Integrons. PLoS Genet 6, e1000793 (2010).

35. Lévesque, C., Brassard, S., Lapointe, J. & Roy, P. H. Diversity and relative strength of tandem promoters for the antibiotic-resistance genes of several integron. Gene 142, 49–54 (1994).

36. Krin, E. et al. Expansion of the SOS regulon of Vibrio cholerae through extensive transcriptome analysis and experimental validation. BMC Genomics 19, (2018).

37. Slauch, J. M. Interplay between Rho, H-NS, spurious transcription, and Salmonella gene regulation. Proc Natl Acad Sci U S A 119, e2211222119 (2022).

38. Solovyev V, S. A. Automatic Annotation of Microbial Genomes and Metagenomic Sequences. in Metagenomics and its Applications in Agriculture, Biomedicine and Environmental Studies 61–78 (Nova Science Publishers, 2011).

39. LaFleur, T. L., Hossain, A. & Salis, H. M. Automated model-predictive design of synthetic promoters to control transcriptional profiles in bacteria. Nat Commun 13, 5159 (2022).

40. Clemens, J. D., Nair, G. B., Ahmed, T., Qadri, F. & Holmgren, J. Cholera. The Lancet 390, 1539–1549 (2017).

41. Colwell, R. R. & Huq, A. Environmental reservoir of Vibrio cholerae. The causative agent of cholera. Ann N Y Acad Sci 740, 44–54 (1994).

42. Sun, H. et al. Effects of intestinal microbiota on physiological metabolism and pathogenicity of Vibrio. Front Microbiol 13, 3193 (2022).

43. Makino, K. et al. Genome sequence of Vibrio parahaemolyticus: a pathogenic mechanism distinct from that of V cholerae. The Lancet 361, 743–749 (2003).

44. Thompson, F. L., Iida, T. & Swings, J. Biodiversity of vibrios. Microbiol Mol Biol Rev 68, 403– 431 (2004).

45. Rapa, R. A. & Labbate, M. The function of integron-associated gene cassettes in Vibrio species: The tip of the iceberg. Frontiers in Microbiology vol. 4 Preprint at 10.3389/fmicb.2013.00385 (2013).

46. Boucher, Y. et al. Recovery and evolutionary analysis of complete integron gene cassette arrays from Vibrio. BMC Evol Biol 6, (2006).

47. Rowe-Magnus, D. A., Guérout, A.-M. & Mazel, D. Super-integrons. Res Microbiol 150, 641– 651 (1999).

48. Michael, C. A. & Labbate, M. Gene cassette transcription in a large integron-associated array. BMC Genet 11, (2010).

49. Tansirichaiya, S., Rahman, M. A., Antepowicz, A., Mullany, P. & Roberts, A. P. Detection of novel integrons in the metagenome of human saliva. PLoS One 11, (2016).

50. Coleman, N., Tetu, S., Wilson, N. & Holmes, A. An unusual integron in Treponema denticola. Microbiology (N Y) 150, 3524–3526 (2004).

51. Mazel, D. Integrons: agents of bacterial evolution. Nat Rev Microbiol 4, 608–620 (2006).

52. Jacquier, H., Zaoui, C., Sanson-le Pors, M.-J., Mazel, D. & Berçot, B. Translation regulation of integrons gene cassette expression by the *attC* sites. Mol Microbiol 72, 1475–1486 (2009).

53. Escudero*, J. A., Loot*, C., Nivina, A. & Mazel, D. The Integron: Adaptation On Demand. Microbiol Spectr 3, (2015).

54. Warman, E. A., Singh, S. S., Gubieda, A. G. & Grainger, D. C. A non-canonical promoter element drives spurious transcription of horizontally acquired bacterial genes. Nucleic Acids Res 48, 4891–4901 (2020).

55. Loot, C. et al. Differences in integron cassette excision dynamics shape a trade-off between evolvability and genetic capacitance. mBio 8, (2017).

56. Warman, E. A. et al. Widespread divergent transcription from bacterial and archaeal promoters is a consequence of DNA-sequence symmetry. Nat Microbiol 6, 746–756 (2021).

57. Shearwin, K., Callen, B. & Egan, J. Transcriptional interference – a crash course. Trends in Genetics 21, 339–345 (2005).

58. Edgar, R. Gene Expression Omnibus: NCBI gene expression and hybridization array data repository. Nucleic Acids Res 30, 207–210 (2002).

59. Heidelberg, J. F. et al. DNA sequence of both chromosomes of the cholera pathogen Vibrio cholerae. Nature 406, 477–483 (2000).

