## Supplementary Material for "Identification of Promoter Activity in Gene-Less Cassettes from *Vibrionaceae* Superintegrons"

**Supplementary Table S1.** Fragments sequences cloned in pDProm

| Name | Origin species | Sequence (5' → 3') |
|---|---|---|
| *mCherry* | - | GCAGTGAGCGCAACGCAATTTTACTTGTACAGCTCGTCCATGCCGCCGGTGGAGTGGCGGCCCTCGGCGC<br>GTTCGTACTGTTCCACGATGGTGTAGTCCTCGTTGTGGGAGGTGATGTCCAACTTGATGTTGACGTTGTAG<br>GCGCCGGGCAGCTGCACGGGCTTCTTGGCCTTGTAGGTGGTCTTGACCTCAGCGTCGTAGTGGCCGCCGT<br>CCTTCAGCTTCAGCCTCTGCTTGATCTCGCCCTTCAGGGCGCCGTCCTCGGGGTACATCCGCTCGGAGGAG<br>GCCTCCCAGCCCATGGTCTTCTTCTGCATTACGGGGCCGTCGGAGGGGAAGTTGGTGCCGCGCAGCTTCA<br>CCTTGTAGATGAACTCGCCGTCTTGCAGGGAGGAGTCCTGGGTCACGGTCACCACGCCGCCGTCCTCGAA<br>GTTCATCACGCGCTCCCACTTGAAGCCCTCGGGGAAGGACAGCTTCAAGTAGTCGGGGATGTCGGCGGG<br>GTGCTTCACGTAGGCCTTGGAGCCGTACATGAACTGAGGGGACAGGATGTCCCAGGCGAAGGGCAGGGG<br>GCCACCCTTGGTCACCTTCAGCTTGGCGGTCTGGGTGCCCTCGTAGGGGCGGCCCTCGCCCTCGCCCTCG<br>ATCTCGAACTCGTGGCCGTTCACGGAGCCCTCCATGTGCACCTTGAAGCGCATGAACTCCTTGATGATGG<br>CCATGTTATCCTCCTCGCCCTTGCTCACCATGGCTTGCTCCTTAGGAGCAAGCCATGAGTAAAGG |
| $P_{lac}$ *attC gfp* | - | CTCACCATGGCTTGCTCCTTAATGTGAGTTAGCTCACTCATTAGGCACCCCAGGCTTTACACTTTATGCTT<br>CCGGCTCGTATGTTGTGTGGAATTGTGAGCGGATAACAATTTCACACAGGAAACAGCTATGAGTTAACAA<br>ACGCCTCAAGAGGGACTGTCAACGCGTGGCGTTTCCAGTCCCATTGAGCCGCGGTGGTTACGGTTGGTGT<br>GTTTGAGTTTTGTGTTATGCGTTGTCAGCCCCTTAGGCGGGCGTTAGCCAGGAGCAAGCCATGAGTAAAG<br>G |
| $P_{lac}$ *attC mCherry* | - | CTCACCATGGCTTGCTCCTTGGCTAACGCCCGCCTAAGGGGCTGACAACGCATAACACAAAACTCAAACA<br>CACCAACCGTAACCACCGCGGCTCAATGGGACTGGAAACGCCACGCGTTGACAGTCCCTCTTGAGGCGTT<br>TGTTAACTCATAGCTGTTTCCTGTGTGAAATTGTTATCCGCTCACAATTCCACACAACATACGAGCCGGAA<br>GCATAAAGTGTAAAGCCTGGGGTGCCTAATGAGTGAGCTAACTCACATTAGGAGCAAGCCATGAGTAAA<br>GG |
| C1 | *V. cholerae* N16961 | CTCACCATGGCTTGCTCCTTTTAGTTGCCAACGAGAAAAAAGTACCTAAAGCAGAATACTTAGGGCTAAT<br>GTCCGTGTGTTTTTGGCTAGGTTTTCGTATTTCAATTTCTAGTTATTAGGTGTTGTGAAAATTGTTCTTTGG<br>TTCCTTGCGCGATTTGGGCTTTCTGTTTCAGAGGCTGTTAGCTTGTTGAAAGCCTGCTCTGCTGCATTGTC<br>GTTCCAAAAGGTTTGTTTGGTGTTCAAAGTCCGTTTTCTACGGCGTTGTAAATTCCAAGTGGTTTCAGTGA<br>CAGGCGCTTTGAAACTGCGCCTGATTTGAGTTTCTTAATACACGTAAAGCTCGTGTTGGCAAAGCGGCAA<br>TCCACTTCAAATGAAAGTGTTAGGTCTGGCAACTAACAAAGCATTCAAGAGGGATTCACAACGCTTGGCA<br>GTTTTGGTTTGAATCAGCTTTAGTGTTTACTGCACAATAGCTTAGGTTGGGTGGTGGCGTTGTTCACCCCT<br>TAATGCGGCGTTATGTAGGAGCAAGCCATGAGTAAAGG |

| | | |
|---|---|---|
| C2 | *V. cholerae* N16961 | CTCACCATGGCTTGCTCCTTTTATGCTTAATCCATAAAAATCAGCAGGTTGTAATTGTCGATTTCTTTGGC<br>ATTTCATTCAGGTTTTGTCGGCAATTCAATATTGTTTTTCGCTGCCTAATGAAGCGGCAATGTATCTGGTG<br>CTGAAAATTCAGTGTTATTGCCTCATTGAGTTGTCGCGCCCATGCGTGGTGAGTGAGTTTGTCGGAGAGG<br>TTTTCACATTGGCCTAGTTGCTAAATGAAAGCCTGCATTGGTCAGTTCATAAGTAAAAGTTGAGCCAGTT<br>CTAATTCAAGGAAATTCGATGTTTGGTCAATAATTTCAGTGAGTTACTCTTGAGTTGGCTCAAATGTAAA<br>GTGTTGAAGCTTAAGCATAACAAACGCCTCAAGAGGGACTGTCAACGCGTGGCGTTTCCAGTCCCATTGA<br>GCCGCGGTGGTTGCAGTTGTTGTGTTTGAGTTTAGTGTTATGCGTTGTCAGCCCCTTAGGCGGGCGTTATG<br>TAGGAGCAAGCCATGAGTAAAGG |
| C3 | *V. cholerae* N16961 | CTCACCATGGCTTGCTCCTTTTAGTTTCTCTCAGGAAAATCATCAAAAATTCACGCACAAGGTTCTGAAAA<br>TTCAGCCTTTATCGTTCAAGCTGCACAATTTGGTTTTGGGGTTTTTCCTGATGTTGATTTGGTAGAAAGCG<br>TTGAAAGTTCAGCGGTTTCAAAACCGCGGATAACCAACAAGCTTTGATGCTTCAATGTTGTCGGACGTTC<br>AACTTGATCAGTTTGGTTTCATAAAAGGTTGAATCATCGCTGTGATCTGGCTTTCTTTGTGCGGACTCTTA<br>ACGTTTTGGCAGCCAACTTCACAGCTTTTGGCGTTGGACGTGTGTTTGCGTGATGCTTTTGCGTAAAATGA<br>CTTTCAAGAAAATGAGTTTAAAAAGTCATTGTTAAACAATAGGCTACGAAACTAACAAACGCCTCAAAG<br>GGACTGCCAACGCGTGGCGTTTCCAGTCCCATTGAGCCGCAGTGGTTTCGGTTGTTGTGTTTGAGTTTTGT<br>GTAATGCGTTGTCAGCCCCTTAGGCGGGCGTTATGTAGGAGCAAGCCATGAGTAAAGG |
| C4 | *V. cholerae* N16961 | CTCACCATGGCTTGCTCCTTTTAGCTTCTTGAAGCAAAAATTATCAAAGTCTGGATTTAATTGTCCTGAAA<br>ATTCAGCTTTTAGTGTTCAAGTTTCACAATTTGGCTTTCGAGTCTTTCTGATGCTTATTTGGTAGAAAGTGT<br>GGAATCTTCAGCAGTTTCAAAGCCGCTAAAAATCATCAAACCTTGATGCTTCAATGTTGTTGGATGCTCC<br>AAACGGAAAGCTTGGTTTCACAAAGGCTGCATCATCGCTGAAAGAGTGTTTTCTTTGTCACGGACTCTTA<br>ACGTGGCCAACTTAACCTTGAATGCTTTGAGTTTTGGCTGTTTACTTCACAGCTTTTGGTGCTGGACGTGC<br>GTTTGTTCGATGCTTTTGCGTAAAATTGCTTTCTCAAAATCATTGTTAAACAATGGGCTACGAAGCTAACA<br>AACGCCTCAAGCGGGACTGTCAACGCGTGGCGTTTCCAGTCCCATTGAGCCGCGGTGGTTTCGATTGTTG<br>AGGTTGAGTTTAGTGTTAATGCGTTGCCAGCCCCTTAGGCGGGCGTTATGTAGGAGCAAGCCATGAGTAA<br>AGG |
| C5 | *V. cholerae* N16961 | CTCACCATGGCTTGCTCCTTTTAGCTTCTTAAAGCAAAAATCGGTTTAATTGTCCTAAAAATTCAGATTTT<br>AGCGTTCAAATTGCACAATTTGGCTTTTGAGCTTTTCTTGATGCTGATTTGGTCGAACGTGTTGAAAGTTC<br>AGTGGGTGCAAAGTCGCTGAAAACTAGTAAGCCTCGATGCTTCAACGTTGTTGGATGTTCCGCTCGGAAA<br>GTTTGATTTCACAAAATGCTGCATCATCGCTGTGATCTTGCTTTCTTTGTCGCGGACTCTTAACCGTGGCC<br>AACTTAACCTTGATCGTTTTAAGTTTTGGTAGCCAACGTCACAGCTTTTGGCGTTGGACGTGTGCTTGCTT<br>GATTCTTTTGCGTAAAATGCCTTTCAAACAAATGTGTTAAAAGCCATTGTTAAACAATAGGCTACGAAGC<br>TAACAAACGCCTCAAGAGGGACTGTCAACGCGTGGCGTTTCCAGTCCCAATGAGCCGCGGTGGTTACGGT |

| | | |
|---|---|---|
| | | TGTTGTGTTTGGGTTTAGTAGTAATGCGTTGCCAGCCCCTTAGGCGGGCGTTATGTAGGAGCAAGCCATG<br>AGTAAAGG |
| C6 | *V. cholerae*<br>N16961 | CTCACCATGGCTTGCTCCTTTTAGCTTCTTAAAGGCTAAGATCATCAAAATCCAAACTTCTTGTTCTGAAA<br>GATCAGTTTTTAGTGTTCAAAGTGCACAATTTGGCATTTCAATTTTGCTAGATGCTGATTTGGTAGAAAGT<br>GTTGGAAGTTCCGCAGTTTCAAAGCCGCTGAAAGCCAACAAGTCTCGATGCTTCAAAGTTGTTGAATGTT<br>CAATTCGGTCAACTTGGTTTCACAAAAAGCAACATCGTCGCTGAAATCGTGCTTCTTTGTCGGAAATGCA<br>GTAAGTGGCCAACTTAGCCTTGAACGCTGCAAGTTTTGGCAGCTCACTTCACAGCTTTTGGCGTTGGACG<br>TGTGTTTGAGTGATGCTTTTGCGTAAAATGCATTTCAAGCAAATGTGTTTAAAAAGTCATTGGTAAACAA<br>TGGGCTACGAAACTAACAAACGCCTCAAGAGGGACTGTCAACGCGTAGCGTTTCCAGTCCCAATGAGCC<br>GTAGTGGTTACGGTTGTTGTGTTTGAGTTTGGTGGAAATGCGTTGCCAGCCCCTTAGGCGGGCGTTATGTA<br>GGAGCAAGCCATGAGTAAAGG |
| C7 | *V. cholerae*<br>N16961 | CTCACCATGGCTTGCTCCTTTTATGCTTAATCCATAAAAATCAGCAGGTTGTGAATGTCGATTCCTTTGGG<br>ATTTCGTTCAGGTTTTGTCGGCAATTCAATATTGTTTGTCGCTGCCTAATGAAGCTGCAATGTGTCTGGCG<br>CTGTAAATTCTGAGTTGTTGCTTCACTAGGTTTTTGAGTCTGCGCGAGGTTAGTCTTGTTGGCTCAAGTCG<br>CTTTGAGCAGTCTGGTGCCTTTACTTACAGGCTTAAAGTCTGTCGGCAATTGAATATTGTTTTGCGCTGCC<br>TAATGAAACAGCAATGTGTCTGTCGCTGTAAATTTCAGTGTTATTGCCTCATTGAGTTGTCGCGCCCATGC<br>GTGGCGAGTGAGTTTGTCGGAGAGGGTTTCCACATTGGCCTAGTTGCTAAATGAAAGCCTGCATTGGTCA<br>GTTCATAGGTGAAATTTTAGCCAGTTCTAATTCAAGGAAACTCGGAATTTGGCCAATGATTTCATTGGGTT<br>ACTCTTGATTTGGTTCAAATGTAAAGTGTTGAAGCTTAAGCATAACAAACGCCTCAAGAGGGACTGTCAA<br>CGCGTAGCGTTTCCAGTCCCATTGAGCCGCGGTGGTTACGGTTGTTGTGTTTGAGTTTGGTGTTATGCGTT<br>GTCAGCCCCTTAGGCGGGCGTTATGTAGGAGCAAGCCATGAGTAAAGG |
| C8 | *V. cholerae*<br>N16961 | CTCACCATGGCTTGCTCCTTTTAGCTTCTTAAAGCAAAAATCATCAAAATCTGGGTTTAATTGTCCTAAAA<br>ATTCAGATTTTAGCGTTCAAATTGCACAATTTGGCTTTTGAGCTTTTCTTGATGCTGATTTGGTCGAACGT<br>GTTGAAAGTTCAGTGGGTGCAAAGTCGCTGAAAACTAGTAAGCCTCGATGCTTCAACGTTGTTGGATGTT<br>CCGCTCGGAAAGTTTGATTTCACAAAATGCTGCATCATCGCTGTGATCTTGCTTTCTTTGTCGCGGACTCT<br>TAACCGTGGCCAACTTAACCTTGATCGTTTTAAGTTTTGGTAGCCAACGTCACAGCTTTTGGCGTTGGACG<br>TGTGCTTGCTTGATTCTTTTGCGTAAAATGCCTTTCAAACAAATGTGTTAAAAGCCATTGTTAAACAATAG<br>GCTACGAAGCTAACAAACGCCTCAAGAGGGACTGTCAACGCGTGGCGTTTCCAGTCCCAATGAGCCGCG<br>GTGGTTACGGTTGTTGTGTTTGGGTTTAGTAGTAATGCGTTGCCAGCCCCTTAGGCGGGCGTTATGTAGGA<br>GCAAGCCATGAGTAAAGG |
| C9 | *V. cholerae*<br>N16961 | CTCACCATGGCTTGCTCCTTTTAGCTTCTTAAAGGCTAAAATCATCAAAATCCCAACTTCTTGTTCTGAAA<br>AATCAGCTTTTAGTATTCAAATTGCACAATTTGGCATTTGAATTTTTCTAGATGCTGATTTGGTAGAAAGT |

| | | |
|---|---|---|
| | | GTTGGAAGTTCCGCAGTTTCAAAGCCGCTGAAAGCCAACAAGCCTCGATGCTTCAAAGTTGTTGGATGTT<br>CAACTCGGTCAACTTGGTTTCACAAAAAGCAGCATTGTCGCTGAAATCGCGCTTTCTTTGTCGGAAATTC<br>GGTAAGTGGCCAACTTAACCTTGATCGCTGCGAGTTTTGGCAGCTCACTTCACAGCTTTTGGCGTTAGAC<br>GTGTGTTTGAGTGATGCTTTTGCGTAAAATGCATTTCAAGCAAATGTGTTTAAAAAGTCATTTTTAAACAA<br>TGGGTTATGAAGCTAACAAACGCCTCAAGAGGGACTGTCAACGCGTGGCGTTTCCAGTCCCATTGAGCCG<br>CGGTGGTTTCGGTTGTTGTGTTTGGGTTTGGTTGTTATGCGTTGTCAGCCCCTTAGGCGGGCGTTATGTAG<br>GAGCAAGCCATGAGTAAAGG |
| C10 | *V. cholerae* N16961 | CTCACCATGGCTTGCTCCTTTTAGTTTCTCTCAATCAAAAATCATCAAAATCTAAGCTCATTTGCATTAAA<br>CATGCAGCTTTTAGTATCCAAATTTCACAAGTTGGCTTTTGAGTTTTGCTTGATGCTGATTTGGCAGAAAG<br>CGTTGAAAACGCAGTAGTTTCAAAGTCGCCGCAAGCCAACAAGTCAAGATGCTTCAACGTTGTTGGGTGT<br>TCAGCGCGGTCAGCTTAGTCTCACAAAAGGCAGCATCATCGCTGTGATCTGGTTTTCTTTGTCGCGGACTA<br>TTAACAGTGGCCAACTTAACCTTGATCGTTTTAAGTTTTGGTAGCTTGCTTCACAGCTTTTGGTGTTGGAC<br>GTGTGCTTGTAAGATGCATTTGCGTAAAATGCTTTTCAAGTAAAGGTGTTAAAAAACCATTGGTAAACAA<br>TGGGCTACGAAACTAACAAACGCCTCAAGAGGGACTGTCAACGCGTGGCGTTTCCAGTCCCATTGAGCC<br>GCGGTGGTTATGGTTGTTGTGTTTGAGTTTAGTGGTAGTGCGTTGCCAGCCCCTTAGGCGGGCGTTATGTA<br>GGAGCAAGCCATGAGTAAAGG |
| F1 | *V. fischeri* ES114 | CTCACCATGGCTTGCTCCTTTTATACGAAATCAATGACGTCCAAATTATAAATATTTAATAGTAACCTACC<br>TGATTTTACAGTTAGTAAGGCAGTGTCAATGTAATAAAATCGTAGTCACTCCTTACTTATGTGTTGCTGTA<br>GACAAGCATTTCATGTCTGGGACTCAATTTTAGCCAATGAGTCCCTTTTTTTTGTGCCGAGGGGGGTAGCG<br>TTAAAAACAAACCTGTCTTATTTTACAGTTGGTTAAGAGTGATTCCAGTAATAAACTGGAGCTCCTCATA<br>ATTTTTGTGCTGCTGTAGACAGGCATTTCACGTTCAGGGTTCGATTCCAGTCATCGAGCCCTTTTTTTATTC<br>ATATCTCTTGTGTTGTTTTAGCTGTGATAAGATGTAATTTATATTAAGTTTTGTATATATGCTTATTCAAAC<br>TAAAGTAGACACGTATAACAAATGCATCAACACGATTTGCTACATTCGGCATTCTAGATTTCTTTGGGTTT<br>GCCGCGTTAAGTGGTAAATTTAAGCTTAATCTGCATGGTAGCAAACGTGTTATGCAGGTGTTATGTAGGA<br>GCAAGCCATGAGTAAAGG |
| F2 | *V. fischeri* ES114 | CTCACCATGGCTTGCTCCTTTTATACGAACTCTGAGAATGCAGTTTTTTGGTTCAGTTCAGCGATTGAACC<br>ATTAGTTTTTTAGGTGTGAAAGTAAGGTGGTTTTCCTTTTTCTTTGTTGCCTTTTAGTGTTCGTTTCATCAG<br>GTTCGGCATTGAACTATTATTGCCCCAAATTCAGGCGTTTCAGAAACTAATTTACGCTTCAATTGTTCATG<br>GTTTTGCCTCATTTTAGCCTGCATCAGTGTTTTCCTCGCAGCATTTTCAATCCCAATTGTGGTTAGGATATT<br>TGGTTTATTAGCTCAGAAGTTGATGAAACCAAATCGTACAAATAGTGTAAGAAAAAGGGATTGTTTCGTT<br>TCACAATTATTGTTGTTGGTTGGATATTTTGCTCGCTTAATTGAGCCAGTCTAATTAAACTCAGTTTACAG<br>CAAATCTTGGTAAATCAGTTCTTTACCAAGTTAATTGGAGCGAAAGCAAGGCTCGTATAACAAAGCAATC |

| | | |
|---|---|---|
| | | AACACGATGCTATTTACATTCGGCATTCGCGGTTTGGAGTATAATTGGTTTTGTAAGTCAGTCTTTCGCAC<br>GTGTTATTGTGGCGTTATGTAGGAGCAAGCCATGAGTAAAGG |
| F3 | *V. fischeri* ES114 | CTCACCATGGCTTGCTCCTTTTGTGCGTACTAAGCCCGAAATTTTTTGCACGGTTCCTCAATAGAAACGAG<br>ACTTTTTTTAAACGCTATTTAGCTATCACAAATCCTCTTCTTTGTTGCCTTTTAGCGTTCGTTTCATCAGGT<br>TCGGCAATTAAGCATTGTCGGTCTAAGTTCAGGCGTTTCAGAGGCATTTTTACGCTTCAATGGTTCTTTGG<br>TGTTGCATCATTTTACTCTGCATCCGTGTTTTCCTCGCAGCATTTTCAATCCCAGTTGTGGCGAGGATATTT<br>GGTTTATCAGCTCAAAATCTGCTGAAACCAAGTCGTTCAAATGGTGTAAAAAATGTATCTGTTTTCCATTT<br>ACTTCATCTCGCAGTTTAAGAACATTTTGTTCGCTTAATTAGCATCGTTAAATTAGAATCAGGGCTTAGAC<br>AATTTTGGCGGGGCAGTTCTTTACCAAGTTGTCCAAGCCGAAAGAGAGTCCGCACAACAATCGCATCAAC<br>ACGATTTACTACACTCGGCAATCTCAGTTTGCCGAGTGTTTCTCGTTTTAGGGCATCAATATTCAGTATAA<br>TTGTATAGTAGTAAACGTGTTATGCAGGCGTTATGTAGGAGCAAGCCATGAGTAAAGG |
| P1 | *V. parahaemolyticus* VPD14 | CTCACCATGGCTTGCTCCTTTTAGGCGACGGGCAGGAAAGTTCTTTGTTCTCATCTTCTTAGTTCTCTTTGA<br>GCATTCGTTTTTCTAAGTCGGCAACTTGGTATTGTCGGCTCAAATTCTTGAATCTCTCCAGCCGTAAAACA<br>TGCCAAATTTCGCGGGTTGCCTCATTGGTTTTCAGTGTTGGATGGTGGTGAGTTTGACTTGCTCAAAGTGT<br>GGCAAGGGCAAGTCAATTGAGGCTTTCGATTGCGCTGCTAGTTTGTCGCGTTGGCATTTTCGGGTTCAAA<br>ACCCGCGTTGTTGTTGAAGGTCGGCATCTCGTGGTCGTTGGAATTTTGGCTGAAACATAGTGAGTTTTGTG<br>GTTCTTGAATCACAAATAATCAGTGTATGTTTGGTTGCTTATTTTGTTTCGGGTAATCTGCTTTCAAACAA<br>CCTTGAAACTGGTGTGAAACTTCGCCTAACAAGGCGTTCAAGACGGATTCACAACGCTTGGCGCTCTCGG<br>TTTTCTTTGAGTTAAGTGATTATGTCACAATGGTTTAGGTAGGGTGGTAGGCGTTGCTCACCACTTAACGC<br>GGCGTTATGTAGGAGCAAGCCATGAGTAAAGG |
| P2 | *V. parahaemolyticus* VPD14 | CTCACCATGGCTTGCTCCTTTTAGTTGCCAACGAGGAAAACGTACCCAAAGCAGATTGTTTAGGGCTAAT<br>GTCCGTGTGTTTTTGGCTAGGTTTTCGTATTTCAATTTCTAGTTATTTAGGTTTGTGAAAATCGATCGTTGG<br>TTTCTCTTGTGGTTGAATTTTCTATTTCAGAAACCGATTTATATGTTGAAAGTTAGCTGTGTAACATTGTCT<br>GTCCAAAGGAGCTTTTTGGTGTTGTTAGCCCGTTTTCTACGGCGTTGTATGTACCAAGTGGTTTCAGTTAC<br>AGTCGCTTTGAATCTGCGCCTGATTTGAGTTTTTCAACGTACGTAAAGTTAGCGTTGGCAAAGTGGCAATT<br>AACTTCAAATCAAAGTGTTAGACTTGGCAACTAACAAAGCATTTAAGAGGGATTCACAACGCTTGGCGGT<br>TTTGGTTTGAATCGGCTTTAGTGTTTACGGCACAATGGTTTAGGTCAGGTGGTGGCGTTGTTCACCCCTTA<br>ATGCGGCGTTATGTAGGAGCAAGCCATGAGTAAAGG |
| P3 | *V. parahaemolyticus* VPD14 | CTCACCATGGCTTGCTCCTTTTAGTCGCCAGAAAAAGCTAGAAGTCAAAATCATGTTTCTAAGCTCAAAA<br>CTTGGCTACTGGTTGTTTGATTCATAGTTCCATCTTCGACGGTCAAAGCTCAGTTTCATCGGTTCTAAATC<br>AATGCAATGCTATTTTCCAAGCGGTTGTTTCGTTCTTTGGTGGTGATAGAGCGGGGCAATCGAAAATCTG<br>TTTTTGGTTGCAAATCTGCTCGGTGAGTTGGTTGGTTTAAAAGCTTATTTACTTGCTGAAATGACGTTTTCT |

| | | |
|---|---|---|
| | | CTGGTGTTGTAAGTTCCACGTGGTTTCAGTGACTTTGTTGAAAGTCCGCACTGTTAGATTGGTTTTCCAAA<br>AGCGCTTTTTGGTGTTGTTAGCCCGTTTTCTGCGGCGCTGTTTTTTCTAAGTGGTTTCATTGAGTAGCCGCT<br>TTGAGACTACGCCTGACTTAAGTGCATCAAAACACATAAAGTTTGTTGTTTGCAAAGTTTAGTTCGGCTTA<br>AAATTGTCGCTCGGTGCCTCGCGACTAACAAACTGCTCAAGAGGGATTCGCAACGCTTGGCATTTTTACT<br>ATGCGTTGAATTTAGTGATTAAGGTGGTATGCGGCGGCTTCGGTATTGCGTTGCTCACCCCTTAGCAGGG<br>CGTTATGTAGGAGCAAGCCATGAGTAAAGG |
| P4 | *V. parahaemolyticus* VPD14 | CTCACCATGGCTTGCTCCTTTTAGGCGACAGGCAGGAAAGTTCTTTGTTCTTACCTTCTTGTTCCCTCTGGC<br>AATTCGCTTTTCGTAGTCGGCAATTAAGCATTGTCGGCTCAACTTCTTGAATCTCTCCAGCCGTAAAACAT<br>GCAAAGGTTCGTGGGTTGCCTCATTAGTTTTCATTGTCGGTTGGTGGATAATTTCACTGGCTCAAAGTGTG<br>GCAGGGGCAAGTTAATCTCGGCTTTCGTTCTCGCTGTAAGTTTGTAGCTTTACGTTTCGCTTGCTCAAAAG<br>CAACTTGAACGCTGAAAGTCAGCATTTTGTGGTTGTGGATATATGGGCTGAAACATAGTGGCTTTTGTGG<br>TTCTTGAACCACAAAAAGTCAGTGGGTGTTTGTTGCTTATAAGGTTTCGGGTAATCTATTTTCAAACCAAC<br>TTAAACACTGGTGGGAAACTTCGCCTAACAAGGCGTTTAAGGCGGATTCACAACGCGTGGCGATTTCAGT<br>TCAAGTCTAGTTTAGTGTTTAAGGTGTAGTGTTTAGGTGTGGTGGTTTGCGTTGTTCACCACTTAACGCGG<br>CGTTATGTAGGAGCAAGCCATGAGTAAAGG |
| V1 | *V. vulnificus* FDAARGOS_663 | CTCACCATGGCTTGCTCCTTTTATGCTTAATCAGGTAAAATCAGTGGTTTATGGTTTTTCTTTGCTCCCTCA<br>GCTTTTCAGTTTGGTGTTTGTCGGCAAGTTGGCTGTTTTGAGCGCTGTTTTTCGGACATCCATTCTTTGGCG<br>CTGGAAATTCCGAGAGTTGCCTCAATCAATTTTTGGGTAAGCGCGAGGTTAGGGCGGTTGGCTCAATCAG<br>AAATCCATTTATAGGTTTTCAAACTTCAAACCAGATTTGCCAAAATTCCTAGTCTGATTATCAAAACTATG<br>ATTCATTTCGGTTTTCTACGTTTTGGTTTTTGTTTGTAAATCAAAGTCGAGTTAATCTTGTTTTTAGTAAAA<br>CGTAAGCCATTGAAGCTTAAGCATAACAAGGCGTTTAAGTGGGATTCATGCCGCGTGGCATTTTGGGTAT<br>GCGGTGAATTTTGGTGGTGAAAGTGGTCTGCGGAAAGTTGGTTTAGGCGGCACTCACCCCTTAACGCGGC<br>GTTATGTAGGAGCAAGCCATGAGTAAAGG |
| V2 | *V. vulnificus* FDAARGOS_663 | CTCACCATGGCTTGCTCCTTTTATGTTTAGTCACGTAAAATCAGTTGGTTGCATCTTTTCTTTGCTTTCTCA<br>GCTTTCTAATTTTGGTCTTGTCAGCAAGTTAACTTCAGTGTTCGCTGTTTTCTGGACTCCAATTCGTGGGCG<br>CGAAAAATTCAGAGAATTGCCTCATTGAAATTCAGTGCGCTTGTGAGTCGAATGAAGCTGCCTTAGTTTT<br>AAGTTTGAGCTCAAGTTTTCTGTTTCTCACGCTAAAACCCAAAGTCATTAAAACTAAATTTATGAGTTAAT<br>TCAGCTTTTTAGCGTCTGGTTTTGGTTTGCAAAGGTAAGTCGAGTTAAGCTCTTGGTATCGCAAACTCAAC<br>CCTTTGTGTCTTAAACATAACAAGGCGTTTAAGCGGGATTCATGCCGCGTGGCATTTTTGGTTTGTAGTGA<br>GTTTTGGTGGTGAAAGTGGTCTGCGGCAGCTTGGTTTATGCGGCATTCACCCCTTAACGCAGCGTTATGTA<br>GGAGCAAGCCATGAGTAAAGG |

| | | |
|---|---|---|
| V3 | *V. vulnificus* FDAARGOS_663 | CTCACCATGGCTTGCTCCTTTTATGTTTAGTCACGTAAAATCAGTTGGTTGCATCTTTTCTTTGTACTCTCG<br>GCTTTCTAATTTTGGTCTTGTCGGCAGGTTAACTTCAGTGTTCGCTGTTTTCTGGACTCCAATTCGTGGGCG<br>CTAAAAATTCAGAGAGTTGCCTCATTAAAGTTCAGTGCGTTTTTGAGTCGAATGAAGCGATCTTAGTTTC<br>AAAGTTTGAGCACCAGTTTTTCGGTTTCTCACGCTAAAACTCAATATCAGTACAATCAAATTTATGAGTTA<br>ATTCAGTTTTTTAGCTTCTAGTTTTGGTTTGCAAAGTTAAGTCGAGTTAAGCTATTGGTTTCGTAAAACTT<br>AACCCTTTGTGTCTTAAACATAACAAGGCGTTTAAGAGGGATTCATGCCGCGTGGCATTTTTGGTATGCG<br>GTTGGTTTTGGTGGTGAAAGTGGTCTGCGGAAGGTTCGTTTATGCGGCATTCACCCCTTAACGCGGCGTT<br>ATGTAGGAGCAAGCCATGAGTAAAGG |
| V4 | *V. vulnificus* FDAARGOS_663 | CTCACCATGGCTTGCTCCTTTTATGCTTAATCACGTAAAATCAGTGGTTTATGGTTTTTCTTTGTTCCCTAG<br>GCTTTTCAGTTTTGTGTTTGTCGGCAAGCCAGCTTTTTTGAGCACTGTTTTTCGGACACTTATTCTTTGGCG<br>CTGGAAATTCAGAGAATTACCTCATTCAATTCTCGGGCAAGCGCGAGGTTAGGGCGGTTGGCTCAATCAG<br>AAATCCATTTTTAGGTTTTCAACTCCAAGCCAGTTTTGCCAAAATCCGAAGTCTGATTATCAAAACTATGA<br>TTCATTTCGGTTTTCTACGTTTTGGTTTTTGTGTGTAAATCAAAGTCGAGTTAATCTTGGTTTTGGTCAAAC<br>GTAACCCATTGAAGCTTAAGCATAACAAGGCGTTTAAGCGGGATTCATGCCGCGTGGCATTTTGGGTTTG<br>CGGTGAGTTTTGGTGGTGAAAGTGGTCTGCAGAAGCTTGGTTTATGCGGCATTCACCCCTTAACGCGGCG<br>TTATGTAGGAGCAAGCCATGAGTAAAGG |
| V5 | *V. vulnificus* FDAARGOS_663 | CTCACCATGGCTTGCTCCTTTTATGTTTAGTCACGTAAAATCAGTTGGTTGCATCTTTTCTTTGTTCCTTCG<br>TATTTCTAATTTTGGTCTTGTCGGCAAGTTAAATTCAGTGGTCGCTGTTTTCTGGACTCCAATTCGTGGGC<br>GCTAAAAATTCAGAGTGTTGCCTCATTTAAGTTCAGTGCGTTTGTGAGTCGAATGAAGCGATGTTAGTTTC<br>AACGTTTGAGCACCAGTTTTCGGTTTCTCACGCTAAAACTTCAATATCAGTACAATCAAAATTATGAGTTA<br>ATTCAGCTTTTTAGCTTCTAACTTTGGTTTTCAAAGTTAAGTCGAGTTAAGCTCTTGGTATCGTAAAACTT<br>AACCCTTTGAGCCTTAAACATAACAAGGCGTTTAAGCGGGATTCATGCCGCGTGGCATTTTTGGTTTGCG<br>TTGAGTTTTGGTGGTGAAAGAGGTCTGCGGAAACTTGGTTTAGGCGGCACTCACCCCTTAACGCAGCGTT<br>ATGTAGGAGCAAGCCATGAGTAAAGG |
| V6 | *V. vulnificus* FDAARGOS_663 | CTCACCATGGCTTGCTCCTTTTATGTTTAGTCACGTAAAAACAGTGGGTTGCATCTATTCTTTGTTTCTTCG<br>TATTTCTAATTTTGGTCTTGTCGGCAAGTTACGTTTAGTGGTCGCTGTTTTCTGGACTCCAATTCGTGGGCG<br>CTAAAAATTCAGAGAGTTGCCTCATTGAATTCCAGAATGTAGGCGAGGTGAGTGAAGTGATTTTGATTTC<br>AAAGTTAGAGCCCCATCTTTTCGATTTCTCACGTTATAAGTCAAAGTCGGCACAATCAAAATTATGATTTA<br>ATTCAGCTTTTTAGCGTCTGGTTTTGGTTTGCAAAGGTAAGTCGAGTTAAGCTATTGGTATCGCAAACTTA<br>ACCCTTTGAGCCTTAAACATAACAAGGCGTTTAAGAGGGATTCATGCCGCGTGGCATTTTTGGTATGCGG<br>TGAGTTTTGGTGGTGAAGGTGGTCTGCGGAAGCTTGGTTTATGCGGCATTCACCCCTTAACGCGGCGTTA<br>TGTAGGAGCAAGCCATGAGTAAAGG |

| | | |
|---|---|---|
| V7 | *V. vulnificus* FDAARGOS_663 | CTCACCATGGCTTGCTCCTTTTATGCTTAATCACGTAAAATCAGTGGTTTATGGCTTTTCTTTGTTCCCTCA<br>GCTTTTTGACTTTGAGCTCGTCGGCAAGTTGGTTTCAGTGGGCGCTGTTTTCTGGACACTTATTCTTTGGC<br>GCTGGAAGTTCAGAGAATTGCCTAAATCAGTTTTCGCGTAAGCGCGAGGTTAGGGCAGTTGGCTTAATCA<br>GATATTTGAACTTGGCTTTTTGAGTTTTAGTCGCCAAATTTCAAAGTCCGATTTTTAAAACAATGAGTCAT<br>TTCGGTTTTCTACGTTTTGTCTTTTGTTTGTAAATCAAAGTCGAGTTAATCTTGGTTCTAGCAAAACGTAAC<br>CCATTGAAGCTTAAGCATAACAAGGCGTTCAAGAGGGATTCATGCCGCGTGGCATTTTTGGTATGCGGCG<br>AGTTTTGGTGGTGAAAGTGGTCTGCTGAAGGTTGGTTTGAGCGGCACTCACCCCTTAACGCAGCGTTATG<br>TAGGAGCAAGCCATGAGTAAAGG |
| V8 | *V. vulnificus* FDAARGOS_663 | CTCACCATGGCTTGCTCCTTTTATGCTTAATCACCTAAAATCAGTGGTTTATGGTTTTTCTTTGTTCCCTAG<br>GCTTTTCAGTTTTGTGTTTGTCGGCAAGCTGGCTTTTTTGAGCGCTGTTTTTCGGACACTTATTCTTTGGCG<br>CTAGAAATTCAGAGAATTGCCTCATTCAATTCTTGGGCAAGCGCGAGGTTAGGGCTGTTGGCTCAATCAG<br>AGATTTGATCTTAGGTTTTTTCGTTTTAGCTGCCAAATTTCAAAGTCTGATTTTCAAAATCATGAGTCATTT<br>CAGTTTTTTGGTTTTTGGCTTTTGTTTGTGAATCAAAGTCGAGTTAATCTTGTTTTTAGTAAAACGTAAGCC<br>ATTGAAGCTTCAGCATAACAAGGCGTTTAAGCGGGATTCATGCCGCGTGGCATTTTTGGTTTGCAGTGAG<br>TTTTGGTGGTGAAAGTGGTGTGCGGAAGGTTGGTTTATGCGGCATTCACCCCTTAACGCAGCGTTATGTA<br>GGAGCAAGCCATGAGTAAAGG |
| V9 | *V. vulnificus* FDAARGOS_663 | CTCACCATGGCTTGCTCCTTTTATGTTTAGTCACGTAAAATCAGTTGGTTGCATCTTTTCTTTGCTTTCTCA<br>GCTTTCTGATTTTGGTCTTGTCGGCAAGTTACGTTTAGTGGTCGCTGTTTTCTGGACTCCAATTCGTGGAC<br>GCTAAAAATTCAAAGAGTTGCCTCATTGAAATTCAGTGCGTTTGTGAGTCGAATGAAGCGATGTTAGTTT<br>CAACGTTTGAGCACCAGTTTTCGGTTTCTCTCGCTAAAATCTCAATATCAGTACAATCAAATTTATGAGTT<br>AATTCAGCTTTTTAGCTTCTAACTTTGGTTTTCAAAGATAAGTCGAGTTAAGCTCTTGGTATTGCAAAACT<br>TACCCCTTTGAGTCTTAAACATAACAAGGCGTTTAAGAGGGATTCATGCCGCGTGGCATTTTTGGTTTGCA<br>GTGAGTTTTGGTGGTGAAAGTGCTCTGCGGAAAGTTGATTTATGCGGCATTCACCCCTTAACGCAGCGTT<br>ATGTAGGAGCAAGCCATGAGTAAAGG |
| V10 | *V. vulnificus* FDAARGOS_663 | CTCACCATGGCTTGCTCCTTTTATGCTTAATCACGTAAAATCAGTGGTTTATGGTTTTTCTTTGTTCCCTCG<br>GCTTTTCATTTTTGTGTTTGTCGGCAAGTTGGCTTTTGTGAGCGCTGTTTTTCGGACACTTATTCTTTGGCG<br>CTGGAAATTCAGAGAATTGCCTCATTCAATTCTTGGGTAAGCGCGAGGTTAGGGCGGTTGGCTCAATCAG<br>AAATCCATTTCTAGGTTTTCAAACTTCAAAATGGATTTGCCAAAATCCCGAACCTGAATATCAAAACTAT<br>GATTCATTTCAGTTTTCTACGTTTTGGTTTTTGTTTGTAAATCAAAGTCGAGTTAATCTTGTTTTTAGTAAA<br>ACGTAAGCCATTGAAGCTTAAGCATAACAAGGCGTTTAAGAGGGATTCATGCCGCGTGGCATTTTTGGTT<br>TGCAGTGAGTTTTGGTGGTGAAAATGGTCTGCGGAAAGTTGGTTTATGCGGCATTCACCCCTTAACGCAG<br>CGTTATGTAGGAGCAAGCCATGAGTAAAGG |

| | | |
|---|---|---|
| V11 | *V. vulnificus*<br>FDAARGOS_663 | CTCACCATGGCTTGCTCCTTTTATGTTTAATCACGTAAAATCAGCCGGTTGCATCTTTTCTTTGTTCCTTCG<br>CATTTCCAATTTTGGTCTTGTCGGCAAGTTGGCTTTCGGTGGTCGCTGGTTTCTGGACTCCAATTCGTGGG<br>CGCGAAAAATTCTGAGAGTTGCCTCATTAAAGCTCAGTGCGTTTGTGAGTCGAATAAAGCGTCTTAGTTT<br>CAAAGTTTGAGCTCCAGCTTTCGGTTTCTCACGCTAAAAATCAAAATCATTAGAATCAAATTTATGAGTT<br>AATTCAGCTTTTTAGCGTCTGATTTTGGTTTTTAAAGGTAACTCGAGTTAAGCTCTTGGTATCGCAAAACT<br>TAACCCTTTGAGCCTTAAACATAACAAGGCGTTTAAGAGGGATTCATGCCGCGTGGCATTTTTAGTATGC<br>GGTGAGTTTTGGTGGTGAAAGTGGTCTGCGGAAACTTGGTTTAGGCGGCACTCACCCCTTAACGCGGCGT<br>TATGTAGGAGCAAGCCATGAGTAAAGG |
| V12 | *V. vulnificus*<br>FDAARGOS_663 | CTCACCATGGCTTGCTCCTTTTATGCTTAATCACGTAAAATCAGTTGGTTATGGTTTTTCTTTGTTCCCTCG<br>GCTTTTCAGTTCGGTGTTTGTCGGCAAGTTGGCTTCTTAGAGCGCTGTTTTTCGGACATTTATTCTTTGGCG<br>CTGGAAATTCAGAGAGTTGCCTCAGTCAATTTTCGGGTAAGCGCTAGGTTAGGGTAGTTGGCTCAATCAG<br>AAATTTGATCTTAGGTTTTTTCGTTTTGGCTGCCACAGTTCAAAGTCTGATTTTCAAAATCATGAGTCATTT<br>CAGTTTTTTGGTTTTTGGCTTTTGTGTGTAAATCAAAGTCGAGTTAATCTTGGTTTTGGTCAAACGTAACC<br>CATTGAAGCTTAAGCATAACAAGGCGTTTAAGAGGGATTCATGCCGCGTGGCATTTTTGGTTTGCAGTGA<br>GTTTTGGTGGTGAAAATGGTCTGCGGAAAGTTGGTTTATGCGGCATTCACCCCTTAACGCAGCGTTATGT<br>AGGAGCAAGCCATGAGTAAAGG |
| V13 | *V. vulnificus*<br>FDAARGOS_663 | CTCACCATGGCTTGCTCCTTTTATGTTTAGTCACGTAAAATCAGTTGGTTGCAGATTTTTTTGCTTTCTAAT<br>TTTGGCCTTGTCGGCAAGCTGGCTTTCGGTGGTCGCTGTTTTCTGGACTCCAATTCGTGGGCGCTAAAAAT<br>TCTGAGAGTTGCCTCATTAAAGCTCAGTGCGTTTGTGAGTCGAATAAAGCGTCTTAGTTTCAAAGTTTGA<br>GCTCTAGTTTTCTGTTTCTCACGTTAAAACTCAAAGTTAGCACAATCAAATTTATGATTTAATTCAGCTTTT<br>TAGCGTCTGGTTTTGGTTTGCAAAGGTAAGTCGAGTTAAGCTATTGGTATCGCAAACTTAACCCTTTGAGC<br>CTTAAACATAACAAGGCGTTTAAGCGGGATTCATGCCGCGTGGCATTTTTTGTTTGCGGTGATTTTTGGTG<br>GTGAAAGTGGTCTGCAGAAGCTTGGTTTATGCGGCATTCACCCCTTAACGCGGCGTTATGTAGGAGCAAG<br>CCATGAGTAAAGG |
| *lacZ*450 | *V. cholerae*<br>N16961 | AAACCCGCACATCGTTAAATGGCACTGCCGTACACCCCATGTTCCTTTGCACAGTTATCGCACTGAGCAG<br>GAGGCTCGTTTGGATGTTGGGGGGAATCGCCAATCTCTAAATGGTCAGTGGCGGTTTGCTCTGTTTGAGA<br>AGCCAGAAGCGGTTGAGCCTGCGGTGATAGACCCGGATTTCGATGATAGCGCTTGGGCGCACATTCCTGT<br>ACCGAGTAACTGGCAGATGCAAGGCTTTGATAAGCCGATTTACACCAATATCCAATATCCATTTGCGGAT<br>CGGCCGCCTTACGTGCCGCAAGATAATCCAACCGGCTGTTATCGCCACCGTTTTACACTGGAAAAACAAG<br>CGCTAACCGAGTCCATTCGCATTGTATTTGATGGGGTCAATTCGGCATTTCATCTGTGGTGCAATGGTCAT<br>TGGGTCGGTTATTCGCAAGATAGCCGCTT |

| | | |
|---|---|---|
| *lacZ*550 | *V. cholerae* N16961 | AAACCCGCACATCGTTAAATGGCACTGCCGTACACCCCATGTTCCTTTGCACAGTTATCGCACTGAGCAG GAGGCTCGTTTGGATGTTGGGGGGAATCGCCAATCTCTAAATGGTCAGTGGCGGTTTGCTCTGTTTGAGA AGCCAGAAGCGGTTGAGCCTGCGGTGATAGACCCGGATTTCGATGATAGCGCTTGGGCGCACATTCCTGT ACCGAGTAACTGGCAGATGCAAGGCTTTGATAAGCCGATTTACACCAATATCCAATATCCATTTGCGGAT CGGCCGCCTTACGTGCCGCAAGATAATCCAACCGGCTGTTATCGCCACCGTTTTACACTGGAAAAACAAG CGCTAACCGAGTCCATTCGCATTGTATTTGATGGGGTCAATTCGGCATTTCATCTGTGGTGCAATGGTCAT TGGGTCGGTTATTCGCAAGATAGCCGCTTGCCTGCCGAGTTTGAGTTAACCCCTTATCTACAAGAGGGTG AAAACCTGTTGGTGGCCATGGTGCTGCGCTGGTCTGATGGCTCTTATTTGGAAGACCAA |
| *lacZ*650 | *V. cholerae* N16961 | AAACCCGCACATCGTTAAATGGCACTGCCGTACACCCCATGTTCCTTTGCACAGTTATCGCACTGAGCAG GAGGCTCGTTTGGATGTTGGGGGGAATCGCCAATCTCTAAATGGTCAGTGGCGGTTTGCTCTGTTTGAGA AGCCAGAAGCGGTTGAGCCTGCGGTGATAGACCCGGATTTCGATGATAGCGCTTGGGCGCACATTCCTGT ACCGAGTAACTGGCAGATGCAAGGCTTTGATAAGCCGATTTACACCAATATCCAATATCCATTTGCGGAT CGGCCGCCTTACGTGCCGCAAGATAATCCAACCGGCTGTTATCGCCACCGTTTTACACTGGAAAAACAAG CGCTAACCGAGTCCATTCGCATTGTATTTGATGGGGTCAATTCGGCATTTCATCTGTGGTGCAATGGTCAT TGGGTCGGTTATTCGCAAGATAGCCGCTTGCCTGCCGAGTTTGAGTTAACCCCTTATCTACAAGAGGGTG AAAACCTGTTGGTGGCCATGGTGCTGCGCTGGTCTGATGGCTCTTATTTGGAAGACCAAGATATGTGGTG GCTGAGTGGCATCTTTCGCGATGTGTATCTCTACCGCAAGCCGATACTCGCGATTGAAGATTTTTTTATCC GCACTGAATTAGATGCGC |

**Supplementary Table S2.** Primers used in this study.

| Oligo | Sequence (5' → 3') | Description |
|---|---|---|
| bb_pSU38_long_F | CTCACCATGGCTTGCTCCTTAGG AGCAAGCCATGAGTAAAGG | Construction of pDProm. Amplification of pA369 backbone. |
| bb_pSU38_long_R | GGACGAGCTGTACAAGTAAAAT TGCGTTGCGCTCACTGC | Construction of pDProm. Amplification of pA369 backbone. |
| bb_pSU38_F | AGGAGCAAGCCATGAGTAAAGG | Amplification of pDProm backbone for cassettes cloning. |
| bb_pSU38_mCherry _R | AAGGAGCAAGCCATGGTGAG | Amplification of pDProm backbone for cassettes cloning. |
| pDProm_lacZ_F | CTCACCATGGCTTGCTCCTTAAA CCCGCACATCGTTAAAT | Amplification of *lacZ* gene. |
| pDProm_lacZ450_R | CCTTTACTCATGGCTTGCTCCTA AGCGGCTATCTTGCGAATAA | Amplification of *lacZ* gene. Construction of pDProm-*lacZ*450. |
| pDProm_lacZ550_R | CCTTTACTCATGGCTTGCTCCTTT GGTCTTCCAAATAAGAGCCA | Amplification of *lacZ* gene. Construction of pDProm-*lacZ*550. |
| pDProm_lacZ650_R | CCTTTACTCATGGCTTGCTCCTG CGCATCTAATTCAGTGCG | Amplification of *lacZ* gene. Construction of pDProm-*lacZ*450. |
| MRVII | GGTTTCCCGACTGGAAAGCG | Check of insert presence in pDProm. |
| MFD | CGCCAGGGTTTTCCCAGTCAC | Check of insert presence in pDProm. |
| bb_pSU38_dfrB9_F | CGTTAGGCGTCGAGCTGCTCTAG ACAGCGCCGTCGTTGC | Construction of pDProm-*drfB9*. Backbone amplification. |
| bb_pSU38_dfrB9_R | TTCGCATTGCGGGGCCTAACAAG GAGCAAGCCATGGTGAG | Construction of pDProm-*drfB9*. Backbone amplification. |
| dfrB9_pSU38_F | CTCACCATGGCTTGCTCCTTGTT AGGCCCCGCAATGCGAA | Construction of pDProm-*drfB9*. Cassette amplification. |
| dfrB9_pSU38_R | GCAACGACGGCGCTGTCTAGAG CAGCGTCGACGCCTAACG | Construction of pDProm-*drfB9*. Cassette amplification. |
| Plac_dfrB9_F | TCACACAGGAAACAGCTATGGT TAGGCCCCGCAATGCGAA | Construction of pDProm $P_{lac}$-*drfB9*. Cassette amplification. |
| bb_pSU38_plac_R | TTCGCATTGCGGGGCCTAACCAT AGCTGTTTCCTGTGTGA | Construction of pDProm $P_{lac}$-*drfB9*. Backbone amplification. |
| bb_dfrB9_E7_F | CACTTAACGCGGCGTTATGTGTT AGGCCCCGCAATGCGAA | Construction of pDProm E7-*drfB9*. Backbone amplification. |

| | | |
|---|---|---|
| E7_dfrB9_R | TTCGCATTGCGGGGCCTAACACA TAACGCCGCGTTAAGTG | Construction of pDProm E7-*drfB9*. Cassette amplification. |
| bb_dfrB9_D11_F | GTGTTATGCAGGCGTTATGTGTT AGGCCCCGCAATGCGAA | Construction of pDProm D11-*drfB9*. Backbone amplification. |
| D11_dfrB9_R | TTCGCATTGCGGGGCCTAACACA TAACGCCTGCATAACAC | Construction of pDProm D11-*drfB9*. Cassette amplification. |
| bb_dfrB9_C12_F | CCCTTAGGCGGGCGTTATGTGTT AGGCCCCGCAATGCGAA | Construction of pDProm C12-*drfB9*. Backbone amplification. |
| C12_dfrB9_R | TTCGCATTGCGGGGCCTAACACA TAACGCCCGCCTAAGGG | Construction of pDProm C12-*drfB9*. Cassette amplification. |
| bb_dfrB9_D1_F | CCCTTAGGCGGGCGTTATGTGTT AGGCCCCGCAATGCGAA | Construction of pDProm D1-*drfB9*. Backbone amplification. |
| D1_dfrB9_R | TTCGCATTGCGGGGCCTAACACA TAACGCCCGCCTAAGGG | Construction of pDProm D1-*drfB9*. Cassette amplification. |
| bb_dfrB9_F11_F | CCCTTAACGCGGCGTTATGTGTT AGGCCCCGCAATGCGAA | Construction of pDProm F11-*drfB9*. Backbone amplification. |
| F11_dfrB9_R | TTCGCATTGCGGGGCCTAACACA TAACGCCGCGTTAAGGG | Construction of pDProm F11-*drfB9*. Cassette amplification. |
| C7_dfrB9_F | GCCTCACCTCGAACGTTATGCTT AATCCATAAAAATCAGC | Construction of pDProm-C8-dfrB9-C7. Cassette amplification. |
| C7_dfrB9_R | GAGCAGCGTCGACGCCTAACGC CCGCCTAAGGG | Construction of pDProm-C8-dfrB9-C7. Cassette amplification. |
| F1_dfrB9_F | TTTGCGCCTCACCTCGAACGTTA TACGAAATCAATGACGTCC | Construction of pDProm-C8-dfrB9-F1. Cassette amplification. |
| F1_dfrB9_R | AGAGCAGCGTCGACGCCTAACA CCTGCATAACACGTTTGC | Construction of pDProm-C8-dfrB9-F1. Cassette amplification. |
| pSU_dfrB9_F | TTAGGCGTCGACGCTGC | Construction of pDProm-C8-dfrB9-F1 and pDProm-C8-dfrB9-C7. Backbone amplification. |
| pSU_dfrB9_R | CGTTCGAGGTGAGGCG | Construction of pDProm-C8-dfrB9-F1 and pDProm-C8-dfrB9-C7. Backbone amplification. |
| gyrA_rt_F | GAGCCAAAGTTACCTTGGCC | *gyrA* amplification for RT-qPCR |

| | | |
|---|---|---|
| gyrA_rt_R | AATGTGCTGGGCAACGACTG | *gyrA* amplification for RT-qPCR |
| gyrA_PRB | [5SUN]-CACCCTCAT-[ZEN]-GGTGACAGTGCGGTTT-[IBFQ] | *gyrA* probe for RT-qPCR |
| gfp_rt_F | GGTGAAGGTGATGCAACATA | *gfp* amplification for RT-qPCR |
| gfp_rt_R | GAAGACCATACGCGAAAGTAG | *gfp* amplification for RT-qPCR |
| gfp_PRB | [6FAM]-ACCTGTTCC-[ZEN]-ATGGCCAACACTTGTCA-[IBFQ] | *gfp* probe for RT-qPCR |
| mCherry_rt_F | CTACTTGAAGCTGTCCTTCC | *mCherry* amplification for RT-qPCR |
| mCherry_rt_R | TAGATGAACTCGCCGTCT | *mCherry* amplification for RT-qPCR |
| mCherry_PRB | [56FAM]-TCAAGTGGG-[ZEN]-AGCGCGTGATGAACT-[IBFQ] | *mCherry* probe for RT-qPCR |
| F1_gsp1_R | GCGGCAAACCCAAAGAAATCTA | 5'RACE of cassette F1 |
| F1_gsp2_R | GAATGCCGAATGTAGCAAATCG | 5'RACE of cassette F1 |
| F1_gsp3_F | CAGTTAGTAAGGCAGTGTCAATGT | 5'RACE of cassette F1 |
| F1_gsp4_R | CATCTTATCACAGCTAAAACAACAC | 5'RACE of cassette F1 |
| F1_ant_gsp1_R | CTCCTTTTATACGAAATCAATGACG | 5'RACE of cassette F1 (antisense) |
| F1_ant_gsp2_R | CAGTTAGTAAGGCAGTGTCAATGT | 5'RACE of cassette F1 (antisense) |
| F1_ant_gsp1_R | TTAACGCGGCAAACCCAAAG | 5'RACE of cassette F1 (antisense) |
| F2_gsp1_R | CCACAATAACACGTGCGAAAGA | 5'RACE of cassette F2 |
| F2_gsp2_R | GACTGACTTACAAAACCAATTATACTC | 5'RACE of cassette F2 |
| F2_gsp3_F | GTGTTCGTTTCATCAGGTTCGG | 5'RACE of cassette F2 |
| F2_gsp4_R | TTATACGAGCCTTGCTTTCGC | 5'RACE of cassette F2 |
| C5_gsp1_R | GGCTGGCAACGCATTACTACT | 5'RACE of cassette C5 |
| C5_gsp2_R | AAACCCAAACACAACAACCGT | 5'RACE of cassette C5 |
| C5_gsp3_F | TGATGCTGATTTGGTCGAACG | 5'RACE of cassette C5 |
| C5_gsp4_R | TCGTAGCCTATTGTTTAACAATGGC | 5'RACE of cassette C5 |
| V11_gsp1_R | GGTGAGTGCCGCCTAAAC | 5'RACE of cassette V11 |
| V11_gsp2_R | CAAGTTTCCGCAGACCACTTTC | 5'RACE of cassette V11 |
| V11_gsp3_F | ATCACGTAAAATCAGCCGGTTG | 5'RACE of cassette V11 |
| V11_gsp4_R | CCAAGAGCTTAACTCGAGTTACC | 5'RACE of cassette V11 |
| V7_gsp1_R | CTCAAACCAACCTTCAGCAGAC | 5'RACE of cassette V7 |
| V7_gsp2_R | CACTTTCACCACCAAAACTCGC | 5'RACE of cassette V7 |
| V7_gsp3_F | GGTTTATGGCTTTTCTTTGTTCCC | 5'RACE of cassette V7 |

| | | |
|---|---|---|
| V7_gsp4_R | TGCTAGAACCAAGATTAACTCG AC | 5’RACE of cassette V7 |
| M13_F | CTGGCCGTCGTTTTAC | Check insert in pTOPO-TA |
| M13_R | GTCATAGCTGTTTCCTG | Check insert in pTOPO-TA |

**Supplementary Table S3.** Putative promoter position and scores predicted by BPROM

| Gene-less cassette or promoter | Strand | Score -10 box | Score -35 box | -10 box | -35 box | LDF |
|---|---|---|---|---|---|---|
| $P_C$S | Sense | 63 | 66 | TCGTAAACT | TTGACA | 5,97 |
| $P_{lac}$ | Sense | 44 | 47 | TCGTATGTT | TTTACA | 3,94 |
| $P_C$W | Sense | 47 | 29 | TCGTAAGCT | TGGACA | 3,10 |
| C1 | Sense | 60 | 40 | TTGTAAATT | TTCAAA | 4,64 |
| | Antisense | 32 | 61 | GTGTATTAA | TTGCCA | 2,37 |
| C2 | Sense | 50 | 38 | AGTTCTAAT | TTCATA | 3,56 |
| | Antisense | 67 | 66 | TGTTATGCT | TTGACA | 6,50 |
| | | 20 | 24 | ATCGACAAT | CTGAAT | 0,91 |
| C3 | Sense | 34 | 36 | AGGTTGAAT | TTGATC | 2,59 |
| | Antisense | 69 | 40 | TTTTAAACT | TTGTTT | 4,15 |
| | | 5 | 38 | TTTCAGAAC | TTGAAC | 0,34 |
| C4 | Sense | 70 | 0 | GCGTAAAAT | GTGCGT | 3,59 |
| | | 32 | 30 | CCTGAAAAT | TTATCA | 1,12 |
| | Antisense | 38 | 66 | TGTTAGCTT | TTGACA | 2,76 |
| | | 24 | 38 | CAGGACAAT | TTGAAC | 1,60 |
| C5 | Sense | 70 | 15 | GCGTAAAAT | GTGCTT | 2,74 |
| | Antisense | 34 | 45 | TTTAACAAT | TTGTTA | 3,61 |
| | | 77 | -6 | CGCTAAAAT | AAGCCA | 3,44 |
| C6 | Sense | 39 | 46 | CGCTGAAAT | TTCACA | 3,22 |
| | Antisense | 42 | 66 | TGTTAGTTT | TTGACA | 2,37 |
| | | 47 | 34 | TAGCAAAAT | TTTCTA | 2,21 |
| C7 | Sense | 56 | 30 | CTGTAAATT | ATGAAA | 3,82 |
| | | 42 | 39 | TGGTGTTAT | TTGTTG | 0,50 |
| | Antisense | 67 | 66 | TGTTATGCT | TTGACA | 6,50 |
| | | -6 | 55 | GAACGAAAT | TTGAAT | 1,04 |
| C8 | Sense | 70 | 15 | GCGTAAAAT | GTGCTT | 2,74 |
| | Antisense | 34 | 45 | TTTAACAAT | TTGTTA | 3,61 |
| | | 77 | -6 | CGCTAAAAT | AAGCCA | 3,44 |
| C9 | Sense | 70 | 7 | GCGTAAAAT | GTGTTT | 2,89 |
| | | 40 | 10 | CTGAAAAAT | ATCAAA | 1,04 |
| | Antisense | 41 | 40 | TTGAATACT | TTCAAA | 2,32 |
| | | 38 | 66 | TGTTAGCTT | TTGACA | 2,28 |
| C10 | Sense | 70 | 15 | GCGTAAAAT | GTGCTT | 4,37 |

| | | | | | | |
|---|---|---|---|---|---|---|
| | Antisense | 38 | 22 | GACTAAGCT | ATGATG | 2,47 |
| P1 | Sense | 60 | 25 | GGGTAATCT | TTGGTT | 5,02 |
| | | 56 | 24 | TGGTATTGT | TTTCTA | 1,84 |
| | Antisense | 56 | 3 | TGACATAAT | CTACCA | 2,93 |
| | | 45 | 61 | AGTCAAACT | TTGCCA | 2,02 |
| P2 | Sense | 59 | 39 | GTGTAACAT | TTTATA | 3,77 |
| | Antisense | 38 | 29 | AGCTAACTT | TGGACA | 4,09 |
| P3 | Sense | 69 | 55 | GCTTAAAAT | TTGCAA | 3,97 |
| | | 54 | 31 | TGCTATTTT | TTTCAT | 1,35 |
| | Antisense | 39 | 55 | TGTTTTGAT | TTGCAA | 3,40 |
| | | 21 | 24 | AAACATGAT | TAGCCA | 0,79 |
| P4 | Sense | 60 | 39 | GGGTAATCT | TTGTTG | 5,54 |
| | Antisense | 37 | 60 | CCTTATAAG | TTGAAA | 4,92 |
| | | 28 | 61 | AGGTAAGAA | TTGCCA | 1,05 |
| V1 | Sense | 57 | 40 | AGTTAATCT | TTGTTT | 4,55 |
| | Antisense | 67 | 25 | TGTTATGCT | ATGAAT | 2,63 |
| | | 13 | 55 | TGAAAAGCT | TTGCCG | 1,01 |
| V2 | Sense | 22 | 60 | GAGTCGAAT | TTGAAA | 2,36 |
| | Antisense | 50 | 20 | AGTTTTAAT | CTGAAT | 2,63 |
| V3 | Sense | 60 | 33 | CAGTACAAT | TTCTCA | 3,21 |
| | Antisense | 57 | 25 | TGTTATGTT | ATGAAT | 3,06 |
| V4 | Sense | 57 | 25 | AGTTAATCT | TTGTGT | 3,87 |
| | Antisense | 67 | 25 | TGTTATGCT | ATGAAT | 2,51 |
| V5 | Sense | 60 | 33 | CAGTACAAT | TTCTCA | 3,14 |
| | Antisense | 57 | 25 | TGTTATGTT | ATGAAT | 3,06 |
| V6 | Sense | 58 | 7 | AATTATGAT | GTCAAA | 4,06 |
| | Antisense | 57 | 25 | TGTTATGTT | ATGAAT | 3,06 |
| V7 | Sense | 57 | 40 | AGTTAATCT | TTGTTT | 4,07 |
| | Antisense | 67 | 25 | TGTTATGCT | ATGAAT | 2,95 |
| V8 | Sense | 57 | 40 | AGTTAATCT | TTGTTT | 4,45 |
| | Antisense | 46 | 38 | ACCTAAGAT | TTGGCA | 3,03 |
| V9 | Sense | 77 | 21 | CGCTAAAAT | TTGAGC | 4,20 |
| | Antisense | 57 | 25 | TGTTATGTT | ATGAAT | 2,96 |
| | | 7 | 55 | CAGAAAGCT | TTGCCG | 0,59 |
| V10 | Sense | 57 | 40 | AGTTAATCT | TTGTTT | 4,66 |
| | Antisense | 67 | 25 | TGTTATGCT | ATGAAT | 2,70 |
| V11 | Sense | 58 | 33 | CATTAGAAT | TTCTCA | 2,64 |

| | | | | | | |
|---|---|---|---|---|---|---|
| | Antisense | 57 | 25 | TGTTATGTT | ATGAAT | 3,06 |
| | | 18 | 32 | AGAAAAGAT | TTGGAA | 0,58 |
| V12 | Sense | 57 | 25 | AGTTAATCT | TTGTGT | 3,99 |
| | Antisense | 67 | 25 | TGTTATGCT | ATGAAT | 2,57 |
| V13 | Sense | 65 | 5 | ATTTATGAT | CTCAAA | 3,35 |
| | Antisense | 57 | 25 | TGTTATGTT | ATGAAT | 3,71 |
| | | 21 | 55 | CAAAAAAAT | TTGCCG | 0,59 |
| F1 | Sense | 61 | 40 | ATTTATATT | TTGTTT | 9,42 |
| | | 46 | 36 | TTTTACAGT | TTTAAT | 2,63 |
| | Antisense | 53 | 55 | ACTTAATAT | TTGAAT | 6,49 |
| | | 76 | 66 | CTGTAAAAT | TTGACA | 6,20 |
| F2 | Sense | 77 | 47 | GAGTATAAT | TTTACA | 6,16 |
| | | 55 | 31 | AACTATTAT | TTTCAT | 3,95 |
| | Antisense | 44 | 36 | GAGCAAAAT | TTTAAT | 4,65 |
| | | 40 | 46 | TGGTTCAAT | TTCACA | 2,81 |
| F3 | Sense | 76 | 27 | CAGTATAAT | TTCTCG | 4,70 |
| | | 31 | 11 | TTTTAAACG | TTCCTC | 2,57 |
| | Antisense | 47 | 52 | CGGCAAACT | TTGATG | 4,71 |
| | | 70 | 17 | GAGTAAAAT | ATGCTG | 2,41 |

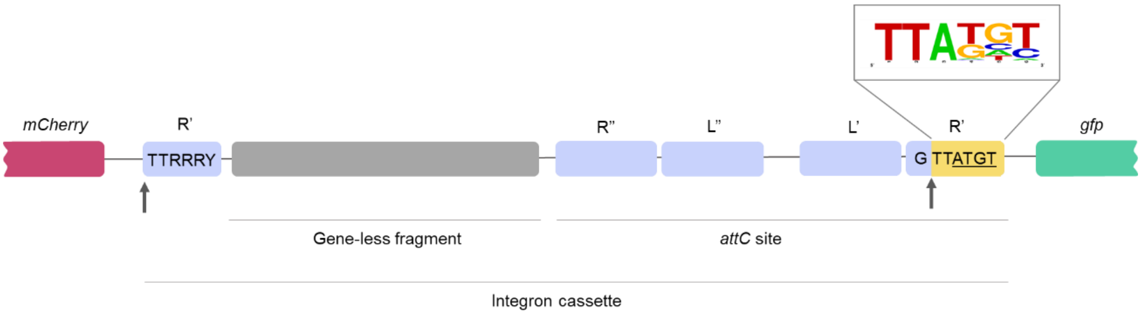


**Supplementary Figure S1. General structure of the synthesized gene-less cassettes cloned in pDProm .** The gene-less DNA fragment for each *Vibrio* spp. was selected including the conserved recombination site R' (G↓TTRRRY) from the recombination point marked with an arrow (TTRRRY), the gene-less fragment, and the *attC* site containing the binding domains R'', L'', L' and R'. After the GTT of the R' recombination point (vertical arrow), the nucleotides ATGT were added to each fragment (yellow box). ATGT were the most prevalent nucleotides from an aligment of all *attC* sites selected in this study (inset).

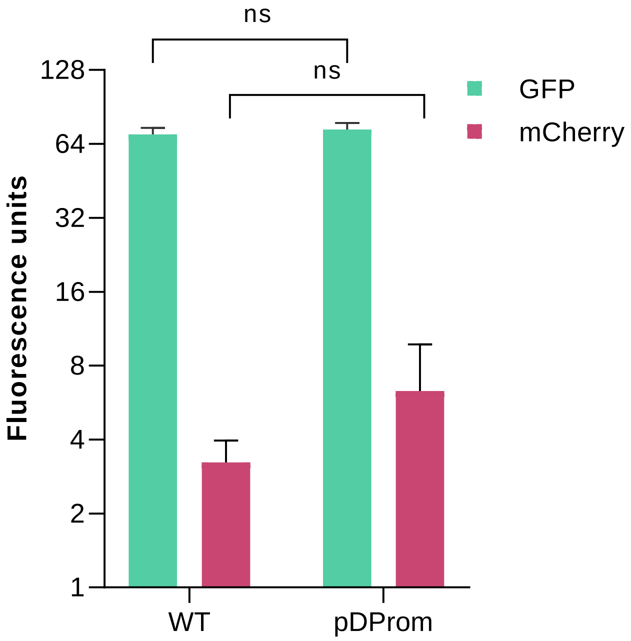


**Supplementary Figure S2. Comparison of fluorescence raw values between *V. cholerae* WT and *V. cholerae* carrying the empty pDProm plasmid.** The fluorescence raw values of both *gfp* (sense) and *mCherry* (antisense) were measured using flow cytometry in *V. cholerae* WT and *V. cholerae* containing the plasmid pDProm without any insert. Error bars represent standard deviation of fluorescence measurements of three biological replicates with two technical replicates each. Paired t-test was performed by comparing the measures obtained for both strains; ns: not significant.

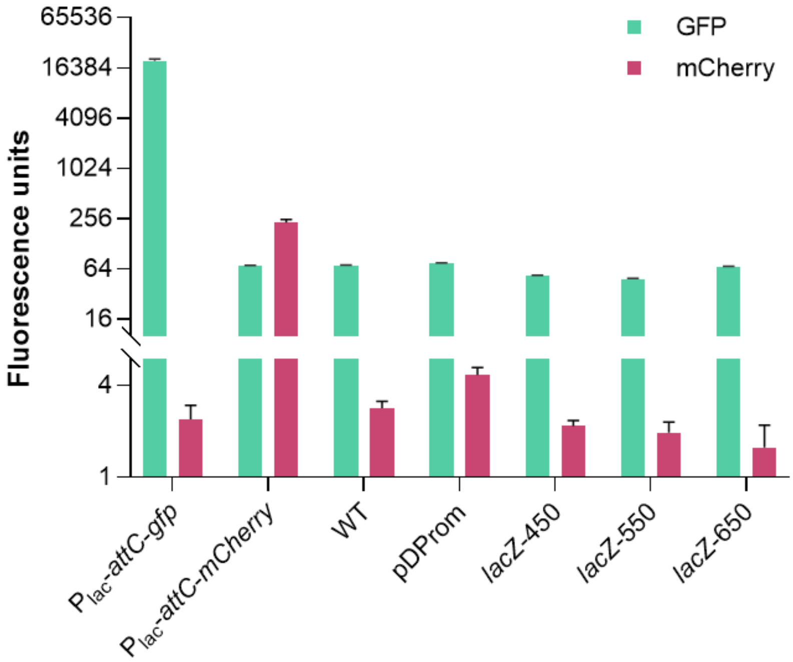


**Supplementary Figure S3. Fluorescence raw values of *V. cholerae* with pDProm and control DNA fragments.** The fluorescence raw values of both *gfp* (sense) and *mCherry* (antisense) were measured using flow cytometry in *V. cholerae* N16961 strains carrying pDProm with control DNA fragments of different sizes (450-, 550-, and 650-bp) originated from the *lacZ* gene. Fluorescence levels were also measured in *V. cholerae* carrying pDProm-$P_{lac}$-*attC-gfp*, pDProm-$P_{lac}$-*attC-mCherry*, the wildtype strain (WT), and the one harboring the empty pDProm. Error bars represent standard deviation of fluorescence measurements of three biological replicates with two technical replicates each.

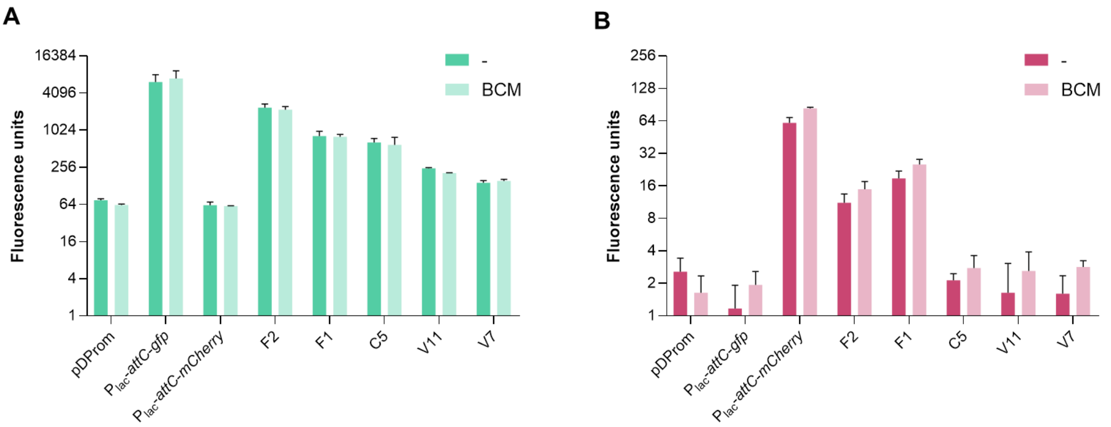


**Supplementary Figure S4. Effect of bicyclomycin (BCM) on fluorescence raw levels of gene-less cassettes.** The fluorescence raw values of both (**A**) *gfp* (sense) and (**B**) *mCherry* (antisense) were measured using flow cytometry in *V. cholerae* strains carrying pDProm and derivatives with or without 2h-incubation with bicyclomycin (50 µg/mL). Error bars represent standard deviation of fluorescence measurements of three technical replicates.

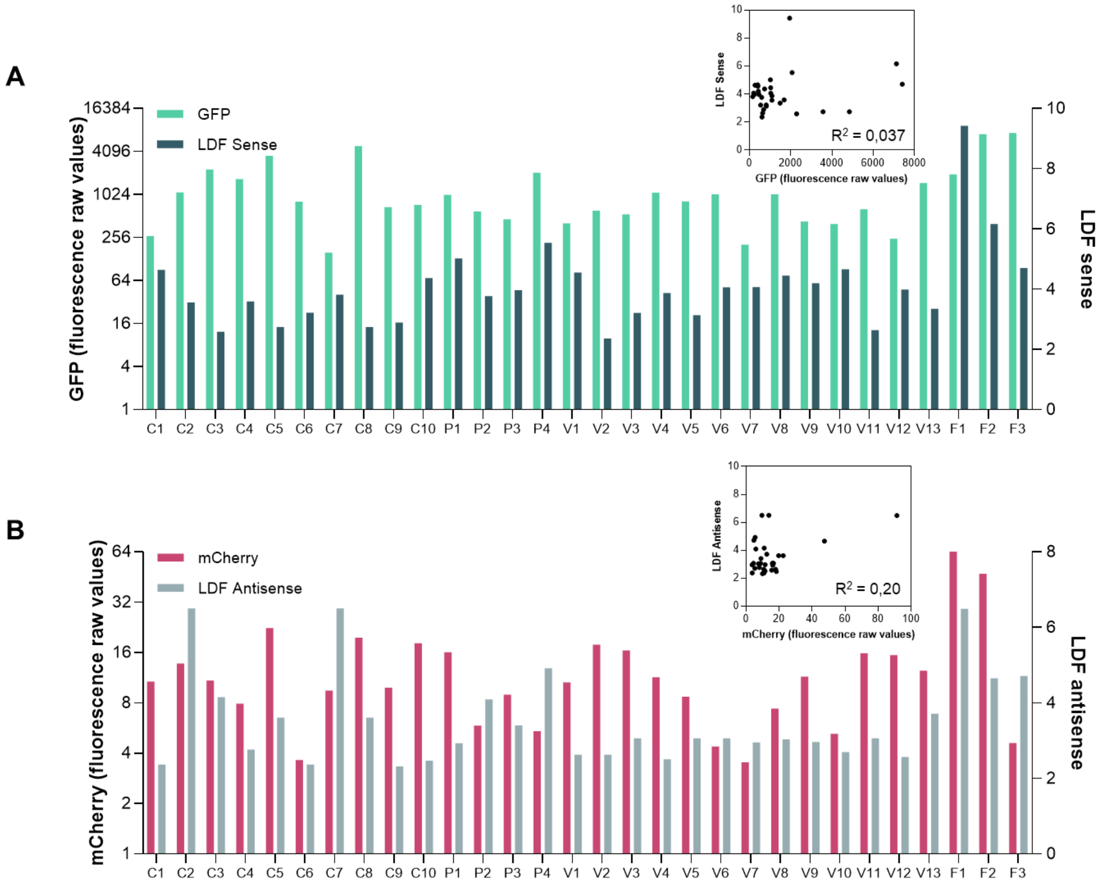


**Supplementary Figure S5. Predicted promoter strength for the *Vibrionaceae* gene-less cassettes compared with the observed expression.** Mean of raw *gfp* fluorescence values (left Y axis) and predicted LDF score of the sense strand by BPROM (right Y axis) of all the selected *Vibrionaceae* cassettes (**A**). Mean of raw *mCherry* fluorescence values (left Y axis) and predicted LDF score of the antisense strand by BPROM (right Y axis) of all the selected *Vibrionaceae* cassettes (**B**). GFP and mCherry fluorescence values do not correlate with the sense and antisense LDF values, respectively (insets: $R^2$=0,037; $R^2$=0,20). The fluorescence mean is calculated from measurements of three biological replicates with two technical replicates each. LDF, linear discriminant function.

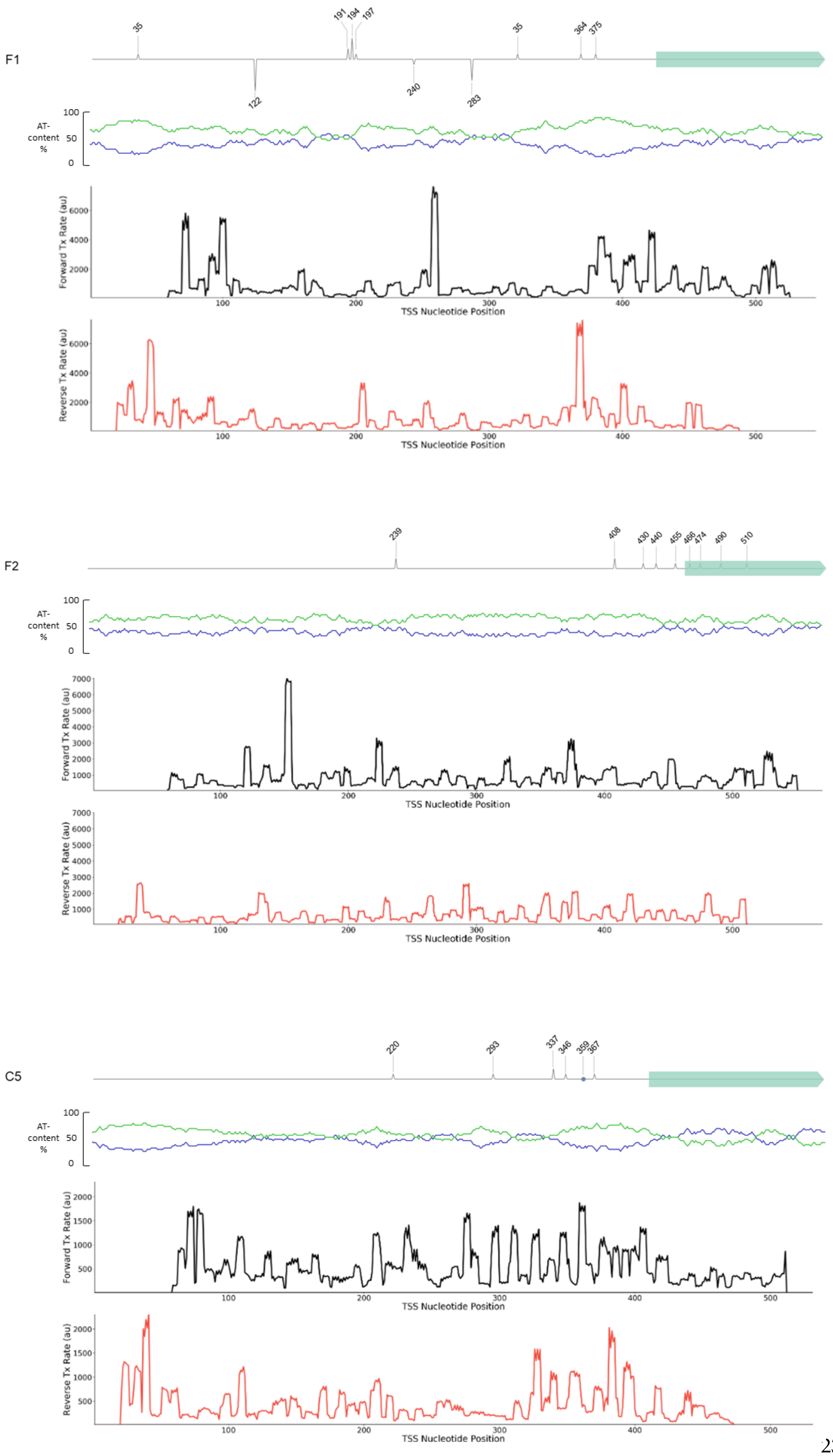
F1
35
191
194
197
35
364
375
122
240
283
AT-content %
Forward Tx Rate (au)
Reverse Tx Rate (au)
TSS Nucleotide Position
F2
239
408
430
440
455
466
474
490
510
AT-content %
Forward Tx Rate (au)
Reverse Tx Rate (au)
TSS Nucleotide Position
C5
220
293
337
346
359
367
AT-content %
Forward Tx Rate (au)
Reverse Tx Rate (au)
TSS Nucleotide Position

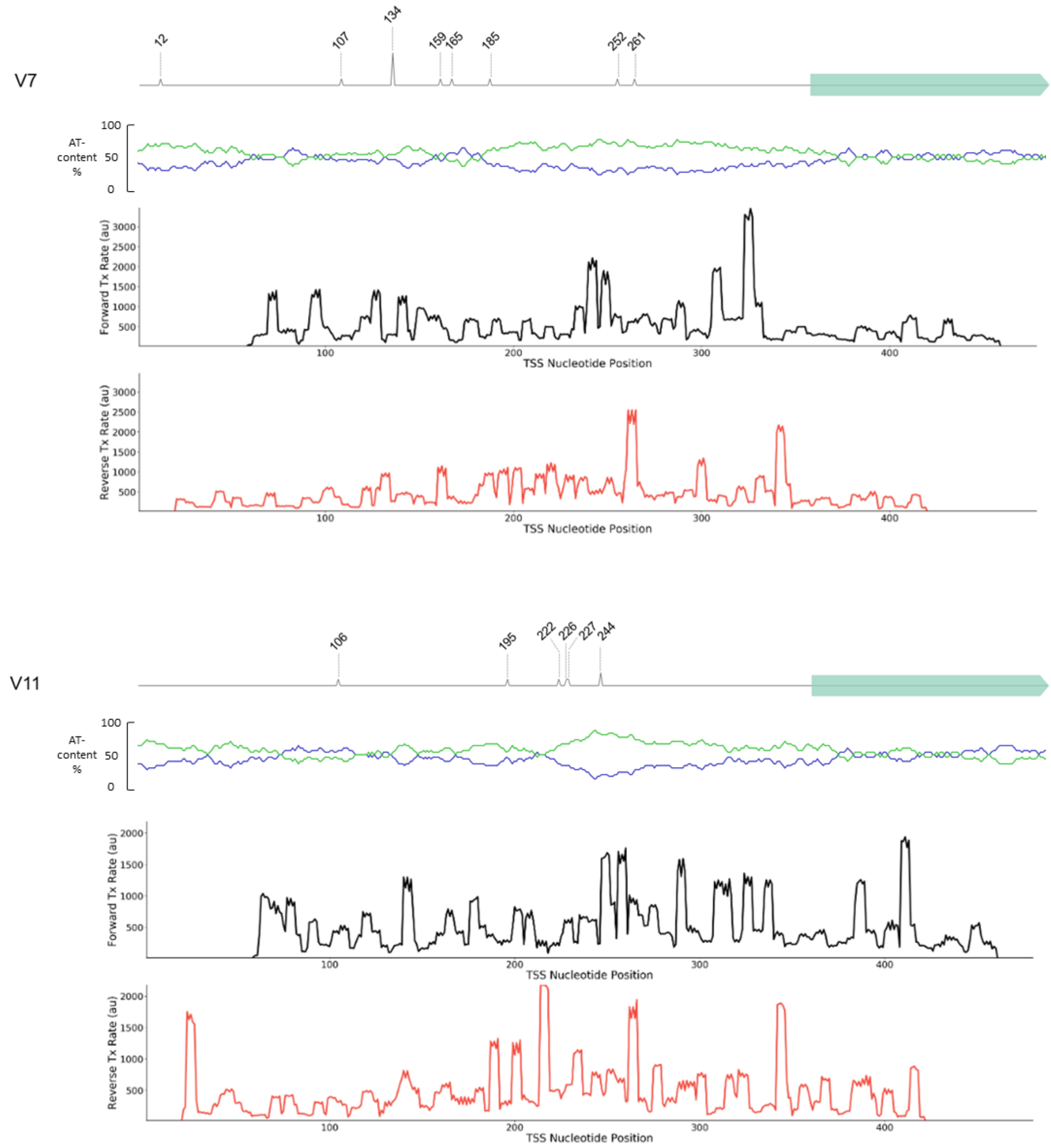


**Supplementary Figure S6. Comparison of detected transcription start sites *in vitro* and *in silico* in gene-less cassettes.** The transcription start sites (TSSs) identified through 5' RACE along the selected cassettes are overlaid with the *in silico* predictions by De Novo DNA software, presented as transcription rates. Graphs illustrating the percentage of AT-content (green curve) and GC-content (blue curve) are also provided for each cassette.

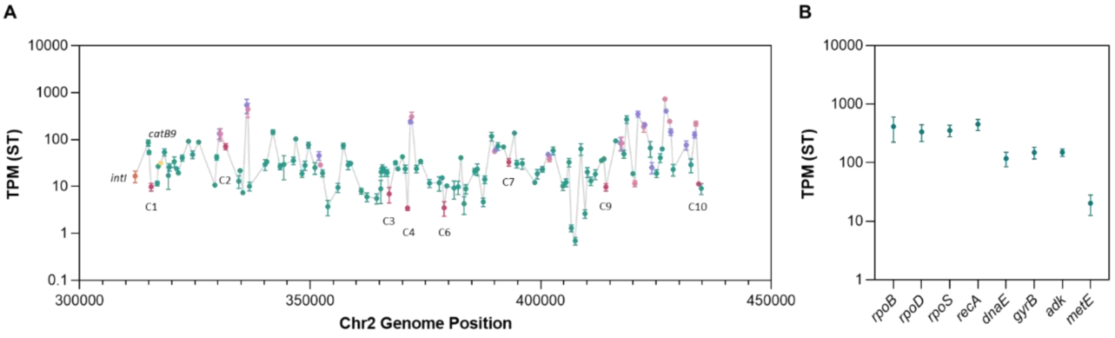


**Supplementary Figure S7. Expression levels of *V. cholerae* N16961 superintegron gene cassettes from BioProject PRJNA420494.** Expression levels represented as transcripts per kilobase million (TPM) of gene cassettes along the superintegron array were calculated from RNA-seq data available in databases in stationary (ST) growth phase (**A**). TPM values of eight housekeeping genes were used as a reference (**B**). TPM values of genes-less cassettes (magenta), toxin-antitoxin systems (mauve and pink), integrase (orange), and the chloramphenicol resistance gene *catB9* (yellow), are highlighted. Error bars represent the standard deviation of three independent biological replicates.

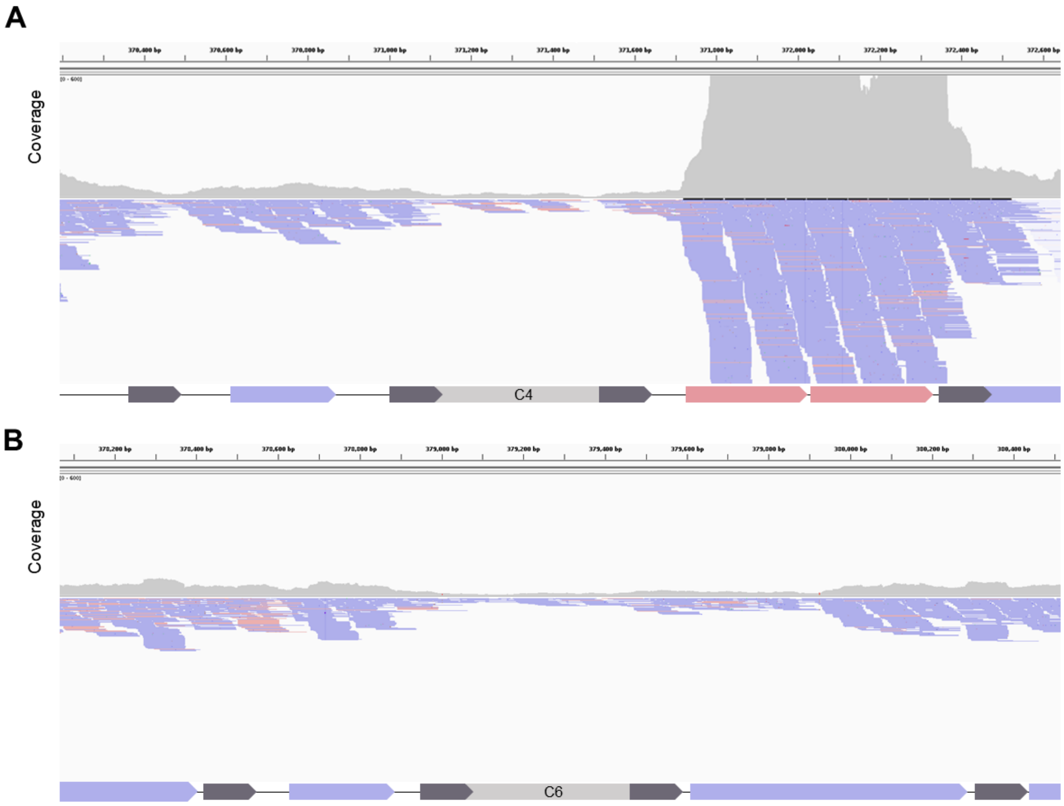


**Supplementary Figure S8. Representation of reads from the RNAseq of *V. cholerae* N16961 superintegron.** The number and orientation of reads, and the coverage upstream and downstream gene-less cassettes C4 (**A**) and C6 (**B**) were analyzed by Integrative Genomics Viewer (IGV) software. Blue color designates genes and reads from the plus (sense) strand; red color designates genes and reads from the minus (antisense) strand.
